# Fusion, fission, and scrambling of the bilaterian genome in Bryozoa

**DOI:** 10.1101/2024.02.15.580425

**Authors:** Thomas D. Lewin, Isabel Jiah-Yih Liao, Mu-En Chen, John D. D. Bishop, Peter W. H. Holland, Yi-Jyun Luo

**Affiliations:** Biodiversity Research Center, Academia Sinica, Taipei, Taiwan; Marine Biological Association, Plymouth, UK; Department of Biology, University of Oxford, Oxford, UK

## Abstract

Groups of orthologous genes are commonly found together on the same chromosome over vast evolutionary distances. This extensive physical gene linkage, known as macrosynteny, is seen between bilaterian phyla as divergent as Chordata, Echinodermata, Mollusca, and Nemertea. Here, we report a unique pattern of genome evolution in Bryozoa, an understudied phylum of colonial invertebrates. Using comparative genomics, we reconstruct the chromosomal evolutionary history of five bryozoans. Multiple ancient chromosome fusions followed by gene mixing led to the near-complete loss of bilaterian linkage groups in the ancestor of extant bryozoans. A second wave of rearrangements, including chromosome fission, then occurred independently in two bryozoan classes, further scrambling bryozoan genomes. We also discover at least five derived chromosomal fusion events shared between bryozoans and brachiopods, supporting the traditional but highly debated Lophophorata hypothesis. Finally, we show that chromosome fusion and fission processes led to the partitioning of genes from bryozoan Hox clusters onto multiple chromosomes. Our findings demonstrate that the canonical bilaterian genome structure has been lost across all studied representatives of an entire phylum; reveal that linkage group fission can occur very frequently in specific lineages; and provide a powerful source of phylogenetic information.

## Introduction

Advances in genome sequencing and assembly have in recent years made it possible to compare the organization of entire genomes across diverse branches of the animal kingdom. The emerging picture is one of a common set of metazoan genes, superimposed with extensive gene duplication, gene loss, and gene novelty in each lineage, plus a surprising degree of large-scale organizational similarity. Many groups of orthologous genes have been retained on the same chromosome for hundreds of millions of years and remain together in diverse and highly divergent lineages (Putnam et al. 2007, 2008; Simakov et al. 2013, 2022; Schultz et al. 2023; Lin et al. 2024). This conservation of chromosome-scale gene linkages, known as macrosynteny, is intriguing, as the selective pressures maintaining it and its significance for genome function are still not fully understood.

The persistence of gene linkages enables the identification of orthologous chromosomes between distantly related phyla. Furthermore, it permits the reconstruction of ancestral linkage groups (ALGs)—hypothesized sets of orthologous genes located on the same chromosome or chromosome arm in ancestral species (Simakov et al. 2022; Mackintosh et al. 2023a). Lineage-specific ALGs have been identified in several animal groups, including vertebrates (Sacerdot et al. 2018; Simakov et al. 2020) and Lepidoptera (butterflies and moths) (Wright et al. 2024). Most strikingly, the discovery of 24 ALGs shared between spiralians and deuterostomes (Simakov et al. 2022; Marlétaz et al. 2023) revealed the genome structure of the most recent bilaterian common ancestor. These highly-conserved bilaterian blocks of orthologous genes are retained in species from phyla as divergent as chordates, echinoderms, molluscs, and annelids (Wang et al. 2017; Simakov et al. 2022; Marlétaz et al. 2023; Martín-Zamora et al. 2023; Lin et al. 2024), suggesting that there are substantial functional constraints to genome organization. There are lineages within phyla that have experienced genomic scrambling, including the breaking up or mixing of genes from different ALGs, such as planarians (Ivankovic et al. 2023), tunicates (Plessy et al. 2024) and clitellate annelids (Schultz et al. 2024; Vargas-Chávez et al. 2024; Lewin et al. 2024a). However, conservation of ALGs seems to be the norm at the phylum level. Despite this general conservation, fluctuations in chromosome number—often leading to ALG rearrangement—are a relatively frequent evolutionary process (White 1954; King 1995; Rice et al. 2015; Román-Palacios et al. 2021). In animals, most of these changes are attributed to inter-chromosomal rearrangements such as translocations and fusions, though they may also arise through whole genome duplication (Orr 1990; Gregory and Mable 2005; Moriyama and Koshiba-Takeuchi 2018; Muffato et al. 2023). Following chromosome fusion, orthologous genes from different ALGs may either be maintained in separate chromosome sections or shuffled and interspersed along the chromosome in a process known as ‘fusion-with-mixing’ (Simakov et al. 2022). Conversely, chromosomes can undergo fission, splitting into separate parts, but the division of ALGs through this process is rare (Simakov et al. 2022; Wright et al. 2024).

The interplay between the stability of orthologous gene sets and the dynamic nature of inter-chromosomal rearrangements shapes the genomic landscape. These rearrangements, while disrupting gene function and expression (Harewood and Fraser 2014) and altering recombination rates (Yoshida et al. 2023), also play a role in driving adaptation (Coyle and Kroll 2007; Dunham et al. 2002; Cheng et al. 2012) and contributing to macroevolutionary processes like speciation (Rieseberg 2001; de Vos et al. 2020; Berdan et al. 2023; Mackintosh et al. 2023b). This dynamic quality of chromosomal changes has made macrosynteny a powerful tool for phylogenetic studies (Steenwyk and King 2024). In particular, fusion-with-mixing events can be particularly powerful phylogenetic markers as the intermixing of genes from different ALGs is considered rare and irreversible. In light of this, macrosynteny was recently used to investigate early branching metazoan lineages (Schultz et al. 2023) and teleost evolution (Parey et al. 2023).

Even in the age of phylogenomics, when the amino acid sequences of thousands of deduced proteins are aligned and analyzed, there remains a small group of ‘Problematica’: taxa for which a robust phylogenetic placement remains tantalizingly out of reach (Jenner and Littlewood 2008; Jenner et al. 2009). Many of these taxa are rare or have few species. Yet one phylum—the Bryozoa—stands out among Problematica as highly abundant, species-rich, and ecologically important (Gordon and Costello 2016; Lombardi et al. 2020), increasingly as alien invasive species (Loxton et al. 2017; Weaver et al. 2018; Pratt et al. 2022). Bryozoans are benthic, suspension-feeding, aquatic invertebrates that are important ecosystem engineers (Wood et al. 2012; Bock and Gordon 2013). They form colonies of individual zooids, generally with calcified skeletons (Zhang et al. 2021), and each zooid can be morphologically differentiated for specialized roles in feeding, brooding, or defense. Despite their unique biological traits, bryozoans and bryozoan genomes remain relatively understudied (Orr et al. 2021; Liao et al. 2023). Several chromosome-scale assemblies have recently been made available (Hoencamp et al. 2021; Bishop et al. 2023a, 2023b; Wood et al. 2023; Bishop et al. 2024), but these are without comparative analysis, and beyond an accepted placement within Spiralia, the phylum’s phylogenetic position remains unresolved (Jenner and Littlewood 2008; Jenner et al. 2009; Khalturin et al. 2022; Liao et al. 2023) (Supplemental Fig. S1).

Here, we present a comparative genomic analysis of five bryozoan genomes, focusing on genome rearrangements and chromosome evolution. Our study aims to (i) characterize the inter-chromosomal rearrangements that have occurred with the phylum Bryozoa; (ii) determine how the 24 bilaterian ALGs are organized in bryozoans; (iii) reconstruct the genome organization of the last common ancestor of extant bryozoans; and (iv) use macrosynteny to elucidate the phylogenetic position of Bryozoa.

## Results

### Chromosome-scale genomes of five bryozoans

To reconstruct the evolutionary history of bryozoan chromosomes, we first assembled a dataset of five chromosome-scale genomes. Four assemblies, *Bugulina stolonifera* (BST) (Wood et al. 2023), *Cryptosula pallasiana* (CPA) (Bishop et al. 2023b), *Membranipora membranacea* (MME) (Bishop et al. 2023a), and *Watersipora subatra* (WSU) (Bishop et al. 2024), are marine bryozoans in the class Gymnolaemata, which accounts for approximately 90% of described bryozoan species (Bock and Gordon 2013) (Fig. 1A). The fifth, *Cristatella mucedo* (CMU) (Hoencamp et al. 2021), is an outgroup belonging to the freshwater class Phylactolaemata (Taylor and Waeschenbach 2015). Four of the five assemblies were generated and made openly accessible by the Darwin Tree of Life (DToL) project (The Darwin Tree of Life Project Consortium et al. 2022), sequenced to chromosome-level using PacBio HiFi long reads and high-throughput chromosome conformation capture (Hi-C) scaffolding (Wood et al. 2023; Bishop et al. 2023a, 2023b, 2024).

**Figure 1.**
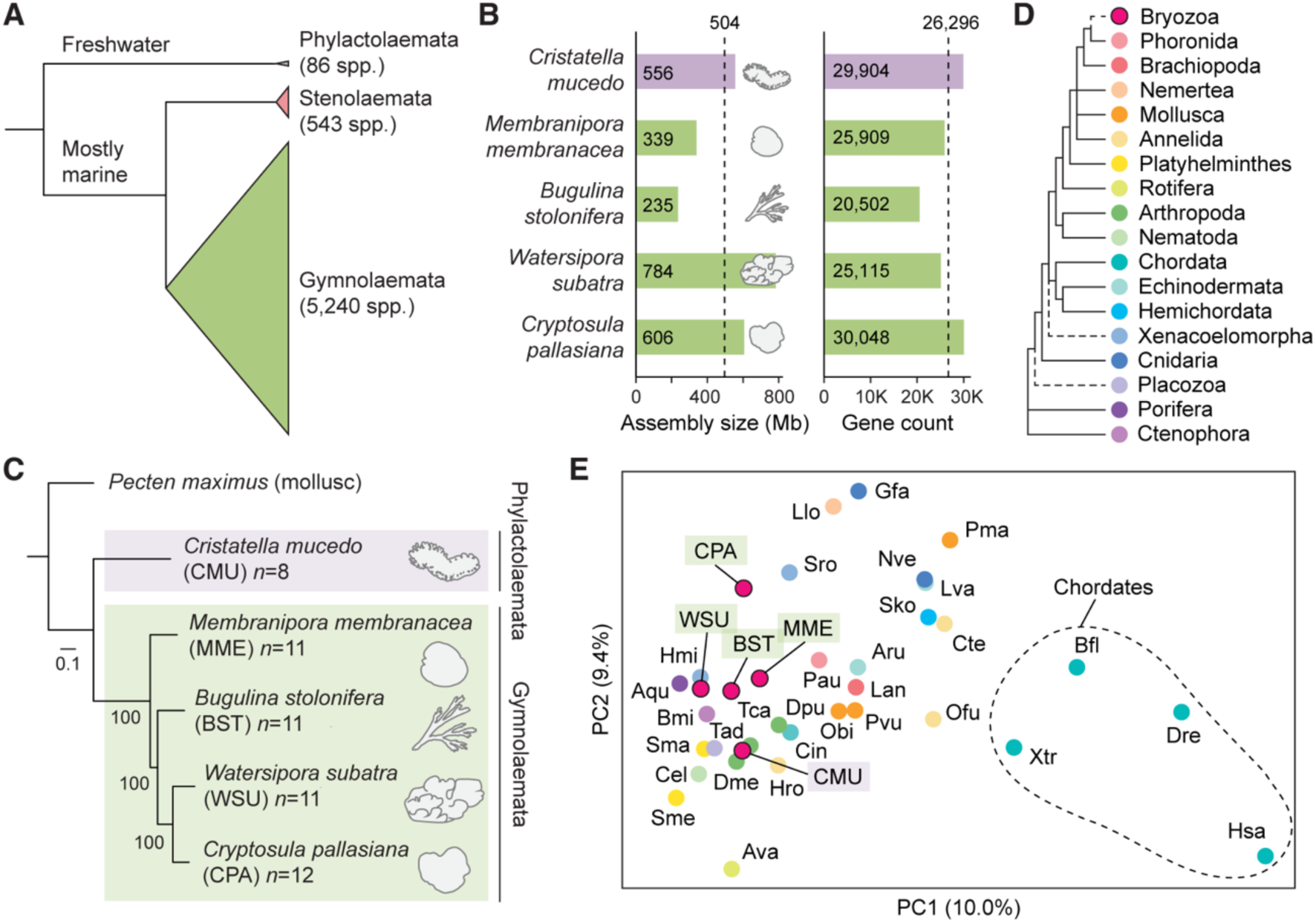
Comparative genomics of five chromosome-scale bryozoan genomes. (*A*) Relationships between the three classes of bryozoans. Class Phylactolaemata is the most basal, containing just 86 species, all of which inhabit freshwater environments (Bock and Gordon 2013). Stenolaemata comprises 543 species, all of which are marine. Gymnolaemata is the most species-rich class, comprising 5,240 species. (*B*) Genome size and gene model count of bryozoan genome assemblies used in this study. Dashed lines mark mean values. The clade containing *W. subatra* and *Cry. pallasiana* has larger genomes. (*C*) Tree of bryozoan species constructed with chromosome-scale genomes with the scallop *Pecten maximus* as the outgroup. The tree was constructed using the maximum likelihood method (LG+F+R6 model) with 1,000 bootstrap replicates in IQ-TREE, utilizing 994 orthologous genes. The scale bar represents the number of substitutions per site. (*D*) Phylogenetic relationships among animal phyla, with dashed lines indicating areas of uncertainty in their placement. Phylogeny based on (Telford et al. 2015) and (Liao et al. 2023). (*E*) Principal component analysis (PCA) conducted using orthologous gene group counts as determined by OrthoFinder. Points are colored by their phylum, as indicated in panel *D*. PCs 1 and 2 explain 10.0% and 9.4% of the variance in the dataset, respectively. Variance ratios for the top 30 principal components are available as Supplemental Fig. S2. Species abbreviations: Ava, *Adineta vaga*; Aqu, *Amphimedon queenslandica*; Aru, *Asterias rubens*; Bfl, *Branchiostoma floridae*; Bmi, *Bolinopsis microptera*; Cel, *Caenorhabditis elegans*; Cin, *Ciona intestinalis*; Cte, *Capitella teleta*; Dme, *Drosophila melanogaster*; Dpu, *Daphnia pulex*; Dre, *Danio rerio*; Gfa, *Galaxea fascicularis*; Hmi, *Hofstenia miamia*; Hro, *Helobdella robusta*; Hsa, *Homo sapiens*, Lan, *Lingula anatina*; Llo, *Lineus longissimus*; Lva, *Lytechinus variegatus*; Nve, *Nematostella vectensis*; Obi, *Octopus bimaculoides*; Ofu, *Owenia fusiformis*; Pau, *Phoronis australis*; Pma, *Pecten maximus*; Pvu, *Patella vulgata*; Sko, *Saccoglossus kowalevskii*; Sma, *Schistosoma mansoni*; Sme, *Schmidtea mediterranea*; Sro, *Symsagittifera roscoffensis*; Tad, *Trichoplax adhaerens*; Tca, *Tribolium castaneum*; Xtr, *Xenopus tropicalis*.

We first performed comprehensive gene prediction using the BRAKER3 pipeline, utilizing transcriptomic data as supporting evidence (Fig. 1B; Supplemental Tables S1–S3). Benchmarking Using Single-Copy Orthologs (BUSCO) (Simão et al. 2015; Manni et al. 2021) reveals a high degree of completeness in gene predictions while also indicating some losses in bryozoans, as evidenced by completeness scores of 82.7% to 88.6% from the Metazoa database (odb10) (Supplemental Table S1). Given accurate phylogenetic relationships are key to inferring chromosome rearrangement events, we built a maximum likelihood phylogenetic tree of the five bryozoan species using gene models obtained from these chromosome-scale genomes (Fig. 1C); this tree is consistent with recent phylogenetic analyses using mitochondrial and ribosomal data (Orr et al. 2022). Principal component analysis of orthologous gene counts finds bryozoans to have a gene set broadly similar to that of other lophotrochozoans (Fig. 1D,E; Supplemental Tables S4,S5).

### Massive chromosome rearrangements within Bryozoa

Haploid chromosome number in our bryozoan dataset ranges from eight in *Cri. mucedo* to 12 in *Cry. pallasiana*. To investigate how the chromosomes of each species are related and understand whether large genome restructuring events have occurred within the phylum Bryozoa, we used OrthoFinder (Emms and Kelly 2019) to identify 4,242 single-copy orthologs and mapped the position of these orthologs within each genome. We then constructed synteny maps which we displayed as ribbon plots (Fig. 2) and Oxford dot plots (Supplemental Fig. S3). Throughout our analysis of chromosome rearrangements, we apply the algebraic symbol conventions established by Simakov et al., wherever applicable (Simakov et al. 2022). Specifically, we use triple bar for one-to-one relationships, also known as syntenic equivalences (≡); tensor for fusion with mixing (⊗); dot product for end-to-end fusion (●); down arrow for centric insertion (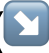); and addition for fission (+) (Fig. 2A). However, although these descriptions are effective for taxa with relatively conserved genomes, we found that the highly derived nature of bryozoan genomes resulted in many chromosomes not fitting clearly into these predefined categories. Furthermore, the notation may become excessively complex and confusing when addressing chromosomes that have undergone numerous ALG fusions and have intricate evolutionary histories. We therefore also utilize narrative descriptions, especially for highly derived chromosomes.

**Figure 2.**
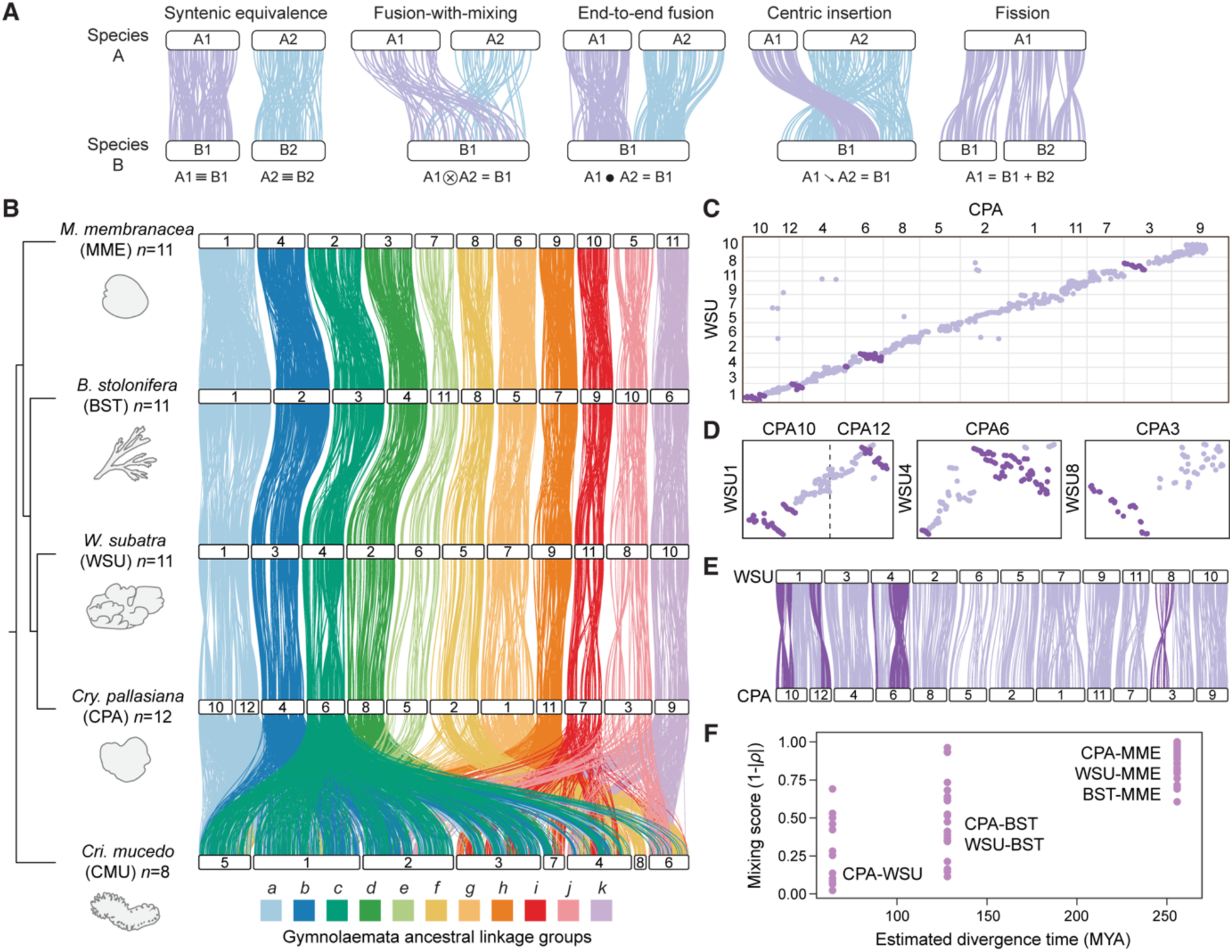
Chromosome rearrangement in gymnolaemate and phylactolaemate bryozoans. *(A)* Schematic representations of possible inter-chromosomal rearrangements. White bars represent chromosomes; colored lines connect orthologous genes; the same color line represents genes in the same ancestral linkage group. (*B*) Chromosome-scale gene linkage between five bryozoan species revealed by ribbon plots. Horizontal bars represent chromosomes. Vertical lines connect the genomic position of orthologous genes in each genome. Chromosomes connected by many orthologs are inferred to be homologous (i.e., orthologous chromosomes). In gymnolaemate species, 11 ALGs *a* to *k* are highly conserved, though the chromosome containing ALG *a* underwent a fission event to form chromosomes CPA10 and CPA12 in *Cry. pallasiana*. These ALGs are not conserved in the phylactolaemate species *Cri. mucedo*. Links between chromosomes sharing five or fewer genes between chromosomes are removed for clarity but can be seen in the Oxford dot plots. (*C*) Oxford dot plot revealing chromosome-scale gene linkage between the two most closely related species in our dataset, *Cry. pallasiana* and *W. subatra*. Each axis represents the entire length of the genome of one species. Gray lines separate chromosomes. Each point represents a pair of orthologs placed by their ordinal position in each genome. Strings of orthologs perpendicular to the main diagonal axis reveal clear intra-chromosomal inversions (dark purple). (*D*) Single-chromosome Oxford dot plots for *Cry. pallasiana* and *W. subatra* chromosomes with clear intra-chromosomal inversions (dark purple). (*E*) Ribbon plot for *Cry. pallasiana* and *W. subatra* highlighting intra-chromosomal inversions (dark purple) (*F*) Microsynteny mixing score versus divergence time for gymnolaemate bryozoan chromosomes. Each point represents a chromosome. Microsynteny mixing score measures how genes are shuffled within a chromosome by intra-chromosomal inversions (see Materials and Methods): a higher score indicates more mixing. Mixing score increases with phylogenetic distance. Abbreviation: MYA, million years ago.

We used the chromosomal position of orthologous genes to identify orthologous chromosomes among the four gymnolaemate species. There is complete syntenic equivalence among three gymnolaemate bryozoans (*B. stolonifera*, *M. membranacea,* and *W. subatra*), which all have a haploid chromosome number of 11, suggesting there have been no instances of chromosome fusion or fission (Fig. 2B). Based on gene position in these species, we assigned 11 gymnolaemate ALGs, labeled ALGs *a* to *k* (Fig. 2B). The final gymnolaemate species, *Cry. pallasiana*, has undergone one fission event to diverge from this state, with the chromosome containing gymnolaemate ALG *a* (*M. membranacea* chr1 ≡ *B. stolonifera* chr1 ≡ *W. subatra* chr1) splitting to form two smaller chromosomes (*Cry. pallasiana* chr10 + chr12) (Fig. 2B). Given the reported rarity of chromosome fission events in animal evolution, we verified that this is not an assembly error by generating a Hi-C contact map, which confirmed that the fission is genuine (Supplemental Fig. S4). This aside, chromosome structure is highly conserved between these species. In contrast, a striking lack of conservation is observed when the phylactolaemate bryozoan *Cri. mucedo* is compared. None of the 11 gymnolaemate ALGs are conserved, and most are completely dispersed and scrambled across the entire *Cri. mucedo* genome (Fig. 2B). For instance, we identified four gymnolaemate ALGs on seven of eight *Cri. mucedo* chromosomes (gymnolaemate ALGs *a*, *b*, *c*, and *g*) (Supplemental Fig. S5). This represents an almost complete loss of chromosome relationships between the two groups. Overall, genome structure is highly conserved within gymnolaemate bryozoans but large inter-chromosomal rearrangements have resulted in highly divergent genome structures between members of Gymnolaemata and Phylactolaemata.

Macrosynteny studies over long evolutionary distances often overlook the precise order of genes, known as microsynteny (Renwick 1971; Passarge et al. 1999). This is because intra-chromosomal rearrangements like inversions rapidly alter gene order, leading to weak microsyntenic signals which are difficult to trace between distantly related species. Within our dataset, there is minimal conservation of gene order between bryozoans with the exception of the two most closely related species, *W. subatra* and *Cry. pallasiana* (Supplemental Fig. S3), which are estimated to have diverged 60–90 million years ago (Orr et al. 2022). Conservation of microsynteny between these species permitted us to investigate the process by which intra-chromosomal rearrangements occur. By generating Oxford dot plots for whole genomes (Fig. 2C) and individual chromosomes (Fig. 2D), we identified six clear cases of large intra-chromosomal inversions between these two species (e.g., *W. subatra* chr1 vs. *Cry. pallasiana* chr12). We noticed many inversions with a complete lack of intermixing, suggesting that these inversions involved entire chromosome arms (e.g., *W. subatra* chr8 vs. *Cry. pallasiana* chr3) (Fig. 2D,E). We developed a microsynteny mixing score (see Methods) to quantify the extent of intra-chromosomal gene shuffling. Between the closely related species *W. subatra* and *Cry. pallasiana*, mixing score varies greatly between chromosomes, suggesting genes are not shuffled at equal rates on different chromosomes (Fig. 2F). In addition, the mixing score increases with estimated divergence time in our dataset (Fig. 2F; Supplemental Tables S6,S7). Overall, the mixing of genes within a chromosome increases with the divergence time between species, and occurs at least in part due to large inversions.

### Near-complete disruption of bilaterian ancestral linkage groups in Bryozoa

To facilitate synteny analysis using bilaterian ancestral linkage groups, we created a new pipeline, SyntenyFinder (see Methods). SyntenyFinder integrates OrthoFinder and RIdeogram and is capable of conducting large-scale comparisons. We first validated our synteny analysis methods using bilaterians with established chromosome relationships. We applied the bilaterian ALG chromosome assignment to the amphioxus *Branchiostoma floridae* and found it to be consistent with previous work (Simakov et al. 2022), identifying bilaterian ALG fusions on chr1 (A1⊗A2), chr2 (C1⊗J2), chr3 (C2●Q), and chr4 (O1●I) . This result confirms the accuracy of our ortholog identification and macrosynteny methodologies.

We next investigated how bryozoan chromosomes relate to the conserved bilaterian genome organization. Using only the bryozoan data, it remains uncertain whether the Gymnolaemata, Phylactolaemata, or neither is similar to the ancestral state of bilaterians. To resolve this, in addition to the amphioxus *B. floridae* (Simakov et al. 2020), we compared the macrosynteny of bryozoans with two other bilaterians with conserved genome organization: the nemertean *Lineus longissimus* (Kwiatkowski et al. 2021) and the mollusc *Pecten maximus* (Kenny et al. 2020). Both *L. longissimus* and *P. maximus* belong to the Lophotrochozoa, the same major protostome clade as bryozoans, while the deuterostome *B. floridae* serves as an outgroup. We generated a dataset of 995 single-copy orthologs and associated each ortholog with one of the 24 bilaterian ALGs described by Simakov et al. (Simakov et al. 2022) (Supplemental Data S1). Synteny maps show that, despite their divergence at the base of the Bilateria, the genome organizations of *B. floridae*, *P. maximus*, and *L. longissimus* exhibit high levels of similarity, having largely preserved the ancestral bilaterian ALGs (Fig. 3; Supplemental Fig. S6; Supplemental Table S8). Our analysis revealed that *P. maximus* and *L. longissimus* share four fusion events that have also been recorded in annelids and brachiopods (J2⊗L, O1⊗R, O2⊗K, and H⊗Q) (Simakov et al. 2022; Martín-Zamora et al. 2023), suggesting that they are ancestral to lophotrochozoans. In addition, we detected several lineage-specific fusion events. Specifically, we identified a fusion on chr2 (B2⊗M) in *P. maximus*; and a fusion on chr2 (C1⊗G) in *L. longissimus*. Overall, genome structures in the deuterostome *B. floridae* and lophotrochozoans *P. maximus* and *L. longissimus* retain most bilaterian ALGs alone on single chromosomes while each have a few lineage-specific fusion events. These changes are modest, considering that these species have been diverging for over half a billion years since the last common ancestor of bilaterians.

**Figure 3.**
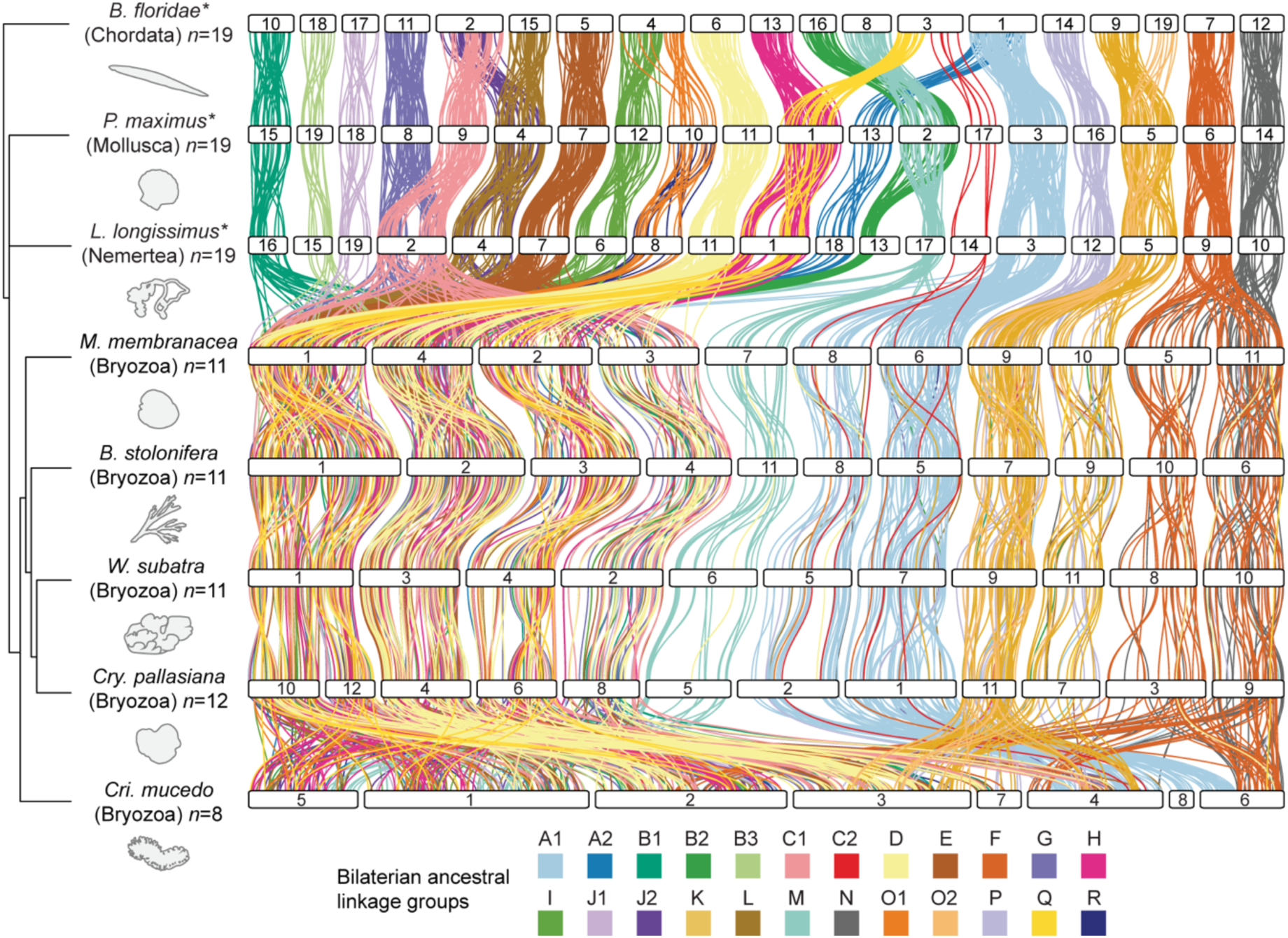
Extensive fusion-with-mixing of bilaterian ancestral linkage groups in bryozoans. Chromosome-scale gene linkage between five bryozoan species and three outgroup bilaterians: a nemertean, a mollusc, and a chordate. Horizontal bars represent chromosomes. Vertical lines connect the genomic position of orthologous genes in each genome. Lines are colored by the bilaterian ALGs defined by (Simakov et al. 2022). These bilaterian ALGs are highly conserved in species ranging from the amphioxus *Branchiostoma floridae* to the scallop *Pecten maximus* and the nemertean *Lineus longissimus*, as indicated by asterisks, but are extensively scrambled in bryozoans.

Like *B. floridae*, *P. maximus*, and *L. longissimus*, bryozoan genomes contain genes from all 24 bilaterian ALGs. However, in contrast to their highly conserved state in other lophotrochozoans and deuterostomes, these bilaterian ALGs have been extensively combined and mixed in the bryozoan lineage (Fig. 3; Supplemental Fig. S6; Supplemental Table S9). We found that 18 out of the 24 bilaterian ALGs (75%) have been combined and mixed across four gymnolaemate chromosomes, indicating that gymnolaemate bryozoans underwent substantial irreversible fusion-with-mixing events followed by fragmentation of a single, ancestrally fused chromosome into four distinct ones. We also detect other bryozoan-specific fusions: ALGs A1 and C2 initially fused (A1⊗C2) and then divided into two chromosomes (e.g., *M. membranacea* chr6 and chr8); ALG P joined with the lophotrochozoan fusion K⊗O2 (P⊗K⊗O2) and later separated into two chromosomes (e.g., *M. membranacea* chr9 and chr10); ALGs F and N fused (F⊗N) and subsequently split into two chromosomes (e.g., *M. membranacea* chr5 and chr11). One chromosome contains only ALG M genes (e.g., *M. membranacea* chr7), although various genes from ALG M were translocated to other chromosomes. The identification of the same chromosome structure across *M. membranacea*, *B. stolonifera*, *W. subatra*, and *Cry. pallasiana* confirms that the observed genome scrambling is not due to assembly errors but rather represents genuine evolutionary divergence. We also found that the *Cri. mucedo* genome arrangement, while unique from other bryozoans, similarly does not preserve the bilaterian ALGs on separate chromosomes. Massive rearrangements have independently taken place within the lineage leading to *Cri. mucedo*, resulting in the dispersion of bilaterian ALGs throughout its genome (Fig. 3). Overall, the genome organizations of the two classes of bryozoans studied here are distinct from each other and highly divergent from the bilaterian ancestral state.

The presence of several ALGs on two or four chromosomes in gymnolaemate bryozoans led us to question whether the splitting of ALGs was due to whole genome duplication (WGD) or chromosome duplications rather than chromosome fission. We tested this using two complementary methods, each using *Metaphire vulgaris*, an annelid with a known WGD (Jin et al. 2020), as a positive control. We first tested for whole genome duplication using synonymous substitutions per site (*K_s_*) distributions for paralogs within each genome. The large peak in the distribution for *M. vulgaris* is caused by its WGD, and the absence of similar large peaks within the *K_s_* distributions of gymnolaemate bryozoans does not support the hypothesis that a recent WGD occurred in this group (Supplemental Fig. S7). There may be very shallow peaks at higher *K_s_* values (*K_s_* = 3 to 4) in the gymnolaemate bryozoans which could indicate ancient WGD in this lineage, but the signal is very weak (Supplemental Fig. S7). Given this uncertainty, we then used MCScanX (Wang et al. 2012) to identify and align collinear blocks of genes within each genome. The presence of duplicated gene blocks on chromosomes with genes from the same ALG would suggest that they were formed by duplication rather than fission. Accordingly, strong pairwise relationships were identified between duplicated chromosomes in *M. vulgaris.* However, in gymnolaemate bryozoans, this analysis revealed zero or few such homologous blocks of genes, which provides no evidence that they are derived from WGD or whole chromosome duplication events (Supplemental Fig. S8). Finally, we used a Duplicate Genes Classifier (Wang et al. 2012) to determine the origin of duplicates found in the genomes. While 23.5% of *M. vulgaris* genes are derived from WGD/segmental duplication events, only 0.4 to 3.6% of genes in the four gymnolaemate species fall into this category (Supplemental Table S10). This again lends no support to the idea of a WGD in gymnolaemate bryozoans. In the phylactolaemate *Cri. mucedo,* we identified several sections of homology between *Cri. mucedo* chromosomes (1 with 2; 1 with 5; 4 with 6) (Supplemental Fig. S8) and several very small very small *K_s_* peaks (Supplemental Fig. S7), suggesting possible segmental duplications in this lineage. Overall, we failed to find evidence that the presence of ALGs on two or four chromosomes in gymnolaemate bryozoans is the product of WGD or large-scale chromosomal duplications, though this cannot be conclusively ruled out.

### The last common ancestor of extant bryozoans had a highly rearranged genome

Since the Gymnolaemata and Phylactolaemata have such distinct genome arrangements, we sought to uncover the genomic structure of their ancestor, the last common ancestor of extant bryozoan lineages. Did it maintain the general bilaterian organization or had rearrangements already occurred? We leveraged the conserved gene linkages found in modern genomes to reconstruct the ancestral state by searching for fusion and fission events that are shared by both gymnolaemate and phylactolaemate species. Since these two classes diverged at the base of the extant Bryozoa, inter-chromosomal rearrangements found in both sets of species can be inferred to have been present in their ancestor. This implicitly assumes that they did not arise multiple times independently, but the complex nature of the shared rearrangements makes that scenario highly improbable. This analysis revealed the likely presence of six linkage groups in the ancestor of living bryozoans, suggesting an ancestral karyotype of six chromosomes (designated here as α, β, γ, δ, ε, and ζ) (Fig. 4A,B). Notably, bryozoan ALGs α and β comprise 10 and eight bilaterian ALGs, respectively, indicating numerous events of sequential fusion-with-mixing (Fig. 4C; Supplemental Fig. S9; Supplemental Table S11). Thus, chromosomal rearrangements had fused and shuffled bilaterian ALGs extensively, even before the extant clades of bryozoans diverged.

**Figure 4.**
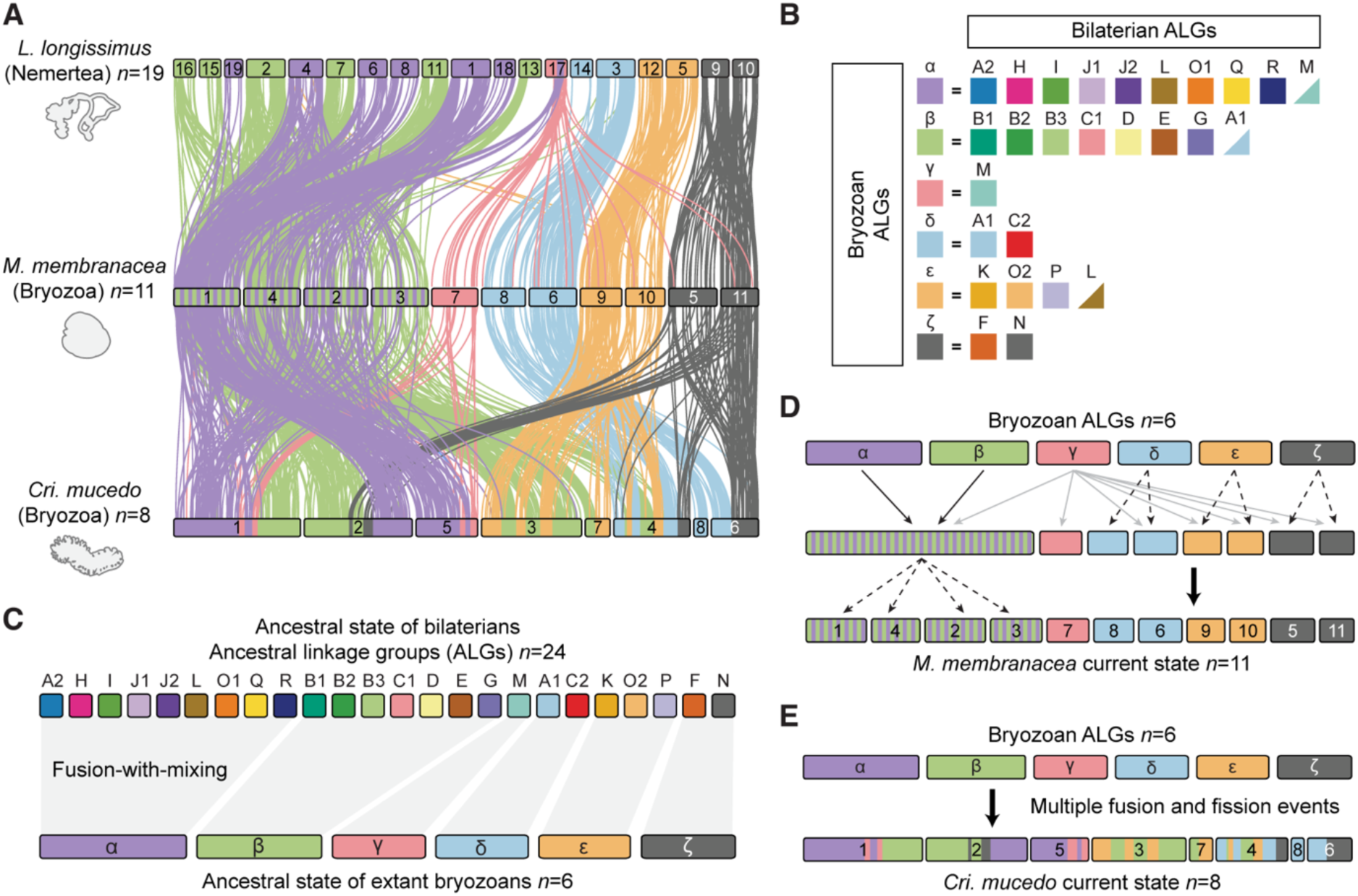
Reconstruction of ancestral genome architecture of extant bryozoans. (*A*) Chromosome-scale gene linkage between *L. longissimus*, *M. membranacea,* and *Cri. mucedo*. Links between orthologs are colored by their 6 bryozoan ALGs. Bryozoan ALGs are inferred from sets of genes that are co-located on the same chromosome (region) in both gymnolaemate and phylactolaemate bryozoans. *L. longissimus* is used as an outgroup because it largely preserves the 24 bryozoan ALGs on separate chromosomes but possess the four fusion events that are shared by lophotrochozoans (J2⊗L, O1⊗R, O2⊗K, and H⊗Q). (*B*) Composition of bryozoan ALGs by bilaterian ALGs, alongside the reconstructed bryozoan ancestral state (six chromosomes: α, β, γ, δ, ε, and ζ). Triangles indicate the inclusion of several genes from the bilaterian ALG but the majority of genes from that ALG are elsewhere. (*C*) Reconstruction of the genome rearrangements that formed the bryozoan ancestral state. The top row shows bilaterian ALGs, and the bottom row shows bryozoan ALGs. (*D*) Reconstruction of the genome rearrangements that led from the bryozoan ancestral state to the extant gymnolaemate genomes. Solid black arrows indicate fusion-with-mixing. Solid gray arrows indicate dispersal around the genome. Dashed arrows indicate chromosome fission. (*E*) Reconstruction of the genome rearrangements that led from the bryozoan ancestral state to the extant phylactolaemate genomes.

Based on this ancestral state of extant bryozoans, we reconstructed the events leading to the observed genome structures in contemporary species (Fig. 4D). In the gymnolaemates, the chromosomes containing ALGs α and β underwent fusion-with-mixing (α⊗β) to create one enormous chromosome containing 18 bilaterian ALGs. This chromosome then underwent sequential fission events to form four daughter chromosomes containing these 18 bilaterian ALGs (e.g., *M. membranacea* chr1 to chr4). Meanwhile, bryozoan ALGs δ, ε, and ζ each underwent a single fission event to form two separate chromosomes. ALG γ was maintained alone on one chromosome (chr7), but genes from this ALG also dispersed to all other chromosomes (Fig. 4D). The *Cri. mucedo* genome exhibits a structure more derived from the ancestral state, characterized by numerous fission and fusion events with limited mixing, resulting in chromosomes consisting of distinct segments derived from different bryozoan ALGs (Fig. 4E). Overall, the bryozoan common ancestor had a highly derived genome, and lineage-specific fusion and fission events have further modified the genomes of both gymnolaemate and phylactolaemate bryozoans.

### Shared derived chromosomal rearrangements between bryozoans and brachiopods

Recent studies have proposed the application of macrosynteny analysis to phylogeny inference, arguing that the infrequency and irreversibility of fusion-with-mixing allows these events to serve as phylogenetic markers (Schultz et al. 2023; Parey et al. 2023; Steenwyk and King 2024). In light of the contentious debate surrounding the phylogenetic position of bryozoans, we sought to determine if macrosynteny could offer insights in this regard. Two contrasting hypotheses exist: (a) the Lophophorata hypothesis, which proposes a close relationship to phoronids and brachiopods (Nesnidal et al. 2013; Luo et al. 2017; Marlétaz et al. 2019; Laumer et al. 2019), and (b) the Polyzoa hypothesis, which proposes closer affiliation to Entoprocta and Cycliophora (Hejnol et al. 2009; Kocot et al. 2017; Khalturin et al. 2022) (Supplemental Fig. S1). Although no chromosome-scale genomes are available for Cycliophora, Entoprocta, or Phoronida species, we recently sequenced a chromosome-level genome for the brachiopod *Lingula anatina* (Lewin et al. 2024b). We first validated the quality of this assembly using genome statistics and found it to be highly contiguous (Supplemental Tables S12,S13), then verified that the chromosome number was supported by cytological karyotype studies (*n* = 10) (Nishizawa et al. 2010). Next, we checked the assembly by comparing chromosome structure to two species with highly conserved genomes: *P. maximus* and *L. longissimus* (Supplemental Fig. S10). The recovery of many ancestral bilaterian ALGs on singular chromosomes further supports that the genome is correctly assembled (Supplemental Fig. S10).

We then conducted a macrosynteny comparison between the bryozoan *M. membranacea*, representative of Gymnolaemata, and the brachiopod *L. anatina* to test for the existence of shared derived rare genomic changes in the form of chromosome fusion-with-mixing events. We identified nine fusion-with-mixing events shared between *L. anatina* and *M. membranacea* (Fig. 5A,B). Four of these events (J2⊗L, O1⊗R, O2⊗K, and H⊗Q) are also shared with annelids, nemerteans, and molluscs (Fig. 3) (Simakov et al. 2022; Martín-Zamora et al. 2023). The remaining five fusion-with-mixing events are exclusive to brachiopods and bryozoans, making them shared derived characteristics unique to these groups (Fig. 5C, marked by ⊗ occurring outside of parentheses). This suggests that bryozoans and brachiopods are closely related, having inherited these changes from a recent common ancestor, supporting the Lophophorata hypothesis (Fig. 5D, scenario Type A). The changes could also be attributed to convergent evolution (Fig. 5E,F, Type B) or a mixture of inheritance from a common ancestor and convergent evolution (Fig. 5G, Type C), but inheritance from a recent common ancestor is the most parsimonious explanation, requiring the fewest fusion-with-mixing events.

**Figure 5.**
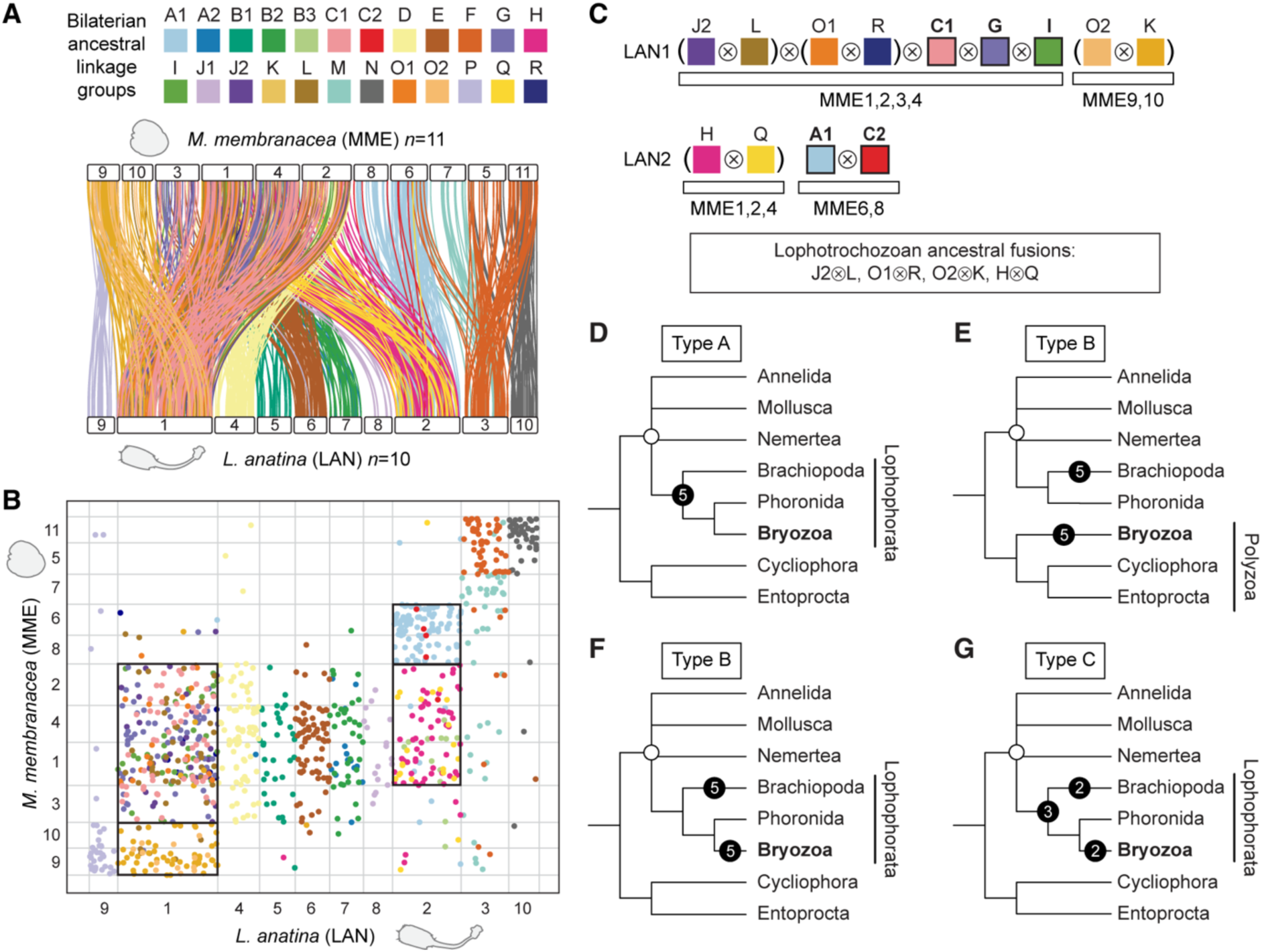
Shared derived chromosome features between bryozoans and brachiopods support the Lophophorata hypothesis. (*A*) Chromosome-scale gene linkage between the bryozoan *M. membranacea* and the brachiopod *L. anatina*. Horizontal bars represent chromosomes. Vertical lines connect the genomic position of orthologous genes in each genome. Lines are colored by bilaterian ALGs. (*B*) Oxford dot plot showing shared ALG fusion events between *L. anatina* and *M. membranacea*. Each point represents a pair of orthologs placed by their ordinal position in each genome, colored by their bilaterian ALG. Black boxes highlight shared fusion-with-mixing events, where two ALGs appear together on the same chromosome in both species. (*C*) There are nine shared fusion-with-mixing events between *L. anatina* and *M. membranacea*. Several spiralian phyla share four of these events (‘lophotrochozoan ancestral fusions’), marked by ⊗ within parentheses, but their timing is uncertain. These events may, therefore, not be useful phylogenetic markers for the position of bryozoans. Confirmed derived fusion-with-mixing events shared between bryozoans and brachiopods but not present in molluscs and annelids are marked by ⊗ occurring outside of parentheses. (*D–G*) Visual representation of four possible evolutionary scenarios that could give rise to the five derived fusion-with-mixing events shared between bryozoans and brachiopods (Types A, B, and C). An open circle marks the base of the Lophotrochozoa. Black circles mark chromosome fusion-with-mixing events. (*D*) Scenario Type A: Brachiopods, phoronids, and bryozoans are closely related, forming the Lophophorata. The five fusion-with-mixing events shared between bryozoans and *L. anatina* occurred in the ancestor of the Lophophorata. Total number of fusions required: 5. (*E*) Scenario Type B-1: Bryozoans, Cycliophora, and Entoprocta are closely related, forming the Polyzoa. Bryozoans are distantly related to brachiopods. The five fusion-with-mixing events shared between bryozoans and *L. anatina* occurred independently in bryozoans and brachiopods. Total number of fusions required: 10. (*F*) Scenario Type B-2: Brachiopods, phoronids, and bryozoans are closely related, forming the Lophophorata. The five fusion-with-mixing events shared between bryozoans and *L. anatina* occurred independently in bryozoans and brachiopods. Total number of fusions required: 10. (*G*) Scenario Type C: Brachiopods, phoronids, and bryozoans are closely related, forming the Lophophorata. Some of the five fusion-with-mixing events shared between bryozoans and *L. anatina* occurred in the ancestor of Lophophorata, while some occurred independently in bryozoans and brachiopods. Total number of fusions required: 6–9.

Under the assumption that fusion-with-mixing events are irreversible, the Type A scenario would be contradicted if any bryozoan, phoronid, or brachiopod were discovered without the described fusion-with-mixing events. To test this, we examined whether the same fusion events are present in *Cri. mucedo*, the most divergent bryozoan in our dataset. Despite *Cri. mucedo* having a highly divergent chromosome organization, all five derived fusion-with-mixing events shared by *M. membranacea* and *L. anatina* also occur together on the same chromosomes in *Cri. mucedo* (Supplemental Fig. S11). Altogether, we observed nine shared fusion chromosome events between bryozoans and brachiopods, with at least five being derived fusion-with-mixing events and thus considered potential phylogenetic markers. These events are notably absent in molluscs, annelids, and nemerteans, which we interpret as indicating a close relationship between bryozoans and brachiopods, supporting the Lophophorata hypothesis. In the future, sequencing a phoronid genome to chromosome level will be a key test of this hypothesis, revealing whether the same fusions are found in this phylum.

### Genome structure and Hox gene evolution

In bilaterians, Hox genes are essential for body patterning and cell type specification (Hubert and Wellik 2023), a role facilitated by their organization into tight clusters (Akam 1989; Graham et al. 1989; Duboule and Dollé 1989). We explored whether extensive genome scrambling has impacted Hox cluster organization in bryozoans. We annotated the Hox genes of the five bryozoan species (Supplemental Fig. S12; Supplemental Data S2) and identified extensive gene loss in the bryozoan Hox gene set, which lacks *Hox1*, *Hox3*, *Scr*, *Antp*, *Lox2*, and *Post1* in all five species. Strikingly, the remaining Hox genes are distributed across two separate chromosomes in every assembly (Fig. 6A). In bilaterians such as *P. maximus* and *L. longissimus*, the Hox cluster is associated with the bilaterian ALG B2. This ALG is fragmented across four chromosomes (five in *Cry. pallasiana*) in each bryozoan genome, as a result of successive fusion and fission events (Fig. 6B). The main Hox cluster is located on orthologous chromosomes containing ALG B2 genes in each bryozoan genome assembly, while Hox genes separated from the cluster are also on a different orthologous chromosome that contains ALG B2 genes (Fig. 6C). This evidence suggests that the presence of Hox genes across two chromosomes in bryozoans is a unique case of partial cluster tandem duplication (*Hox4*, *Lox5*, *Lox4*, and *Post2* genes) followed by separation due to chromosome fission (Fig. 6D).

**Figure 6.**
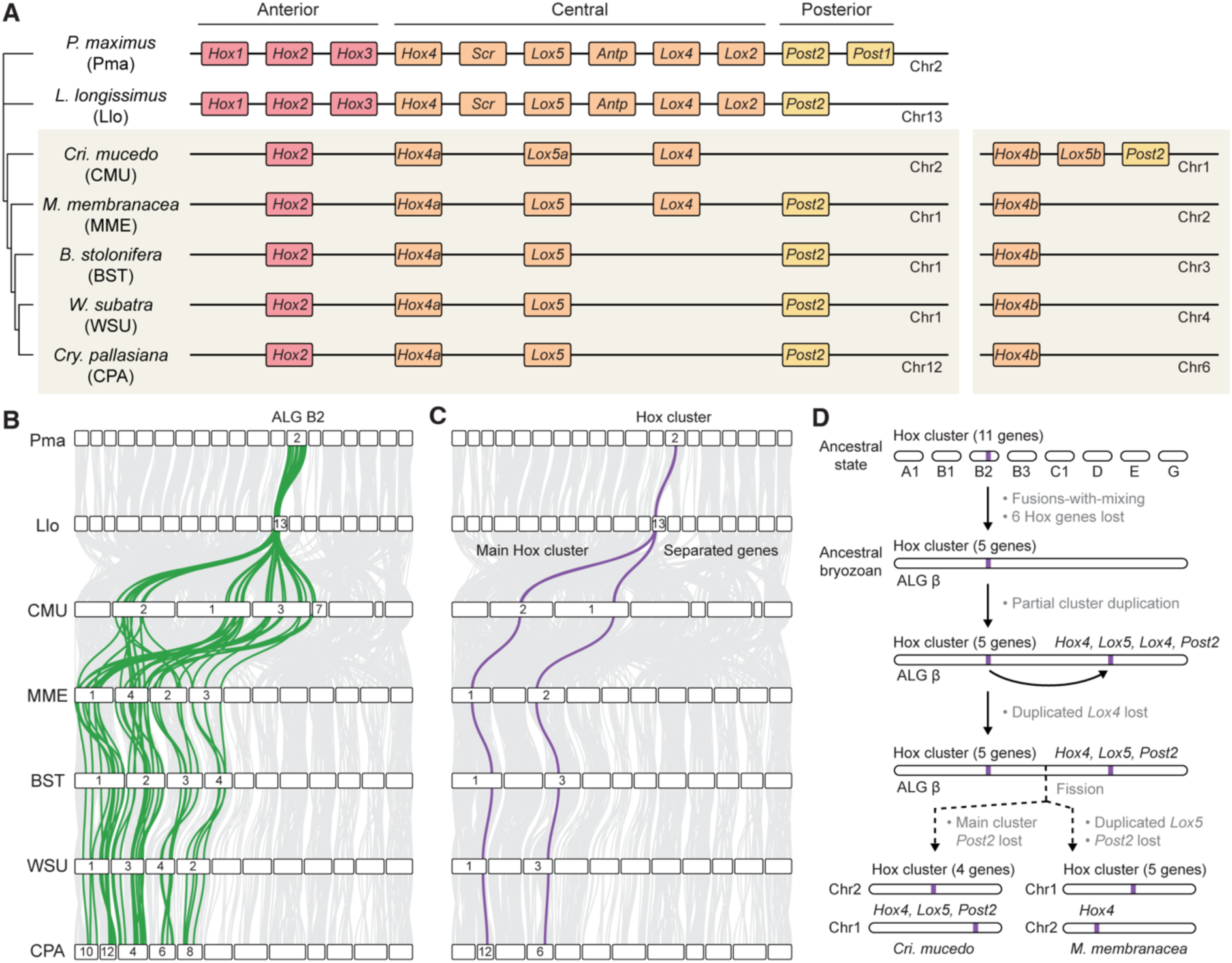
Evolution of the bryozoan Hox cluster. (*A*) Left: Representation of the main Hox cluster of bryozoans and two outgroups, the nemertean *L. longissimus* and the mollusc *P. maximus*. Colored rectangles represent Hox genes. Horizontal black lines represent chromosomes. Right: Separate Hox genes located in different chromosomes. (*B*) Positions of ALG B2 genes (green) in bryozoan genomes. (*C*) Positions of Hox genes (purple) in bryozoan genomes. (*D*) Representation of the evolutionary events that resulted in observed bryozoan Hox gene arrangements. In the bryozoan ancestral lineage, chromosomes containing bilaterian ALGs A1, B1, B2 (including the Hox cluster), B3, C1, D, E, and G underwent sequential fusion events to form bryozoan ALG β; the Hox cluster underwent a partial duplication, copying genes *Hox4*, *Lox5*, *Lox4*, and *Post2*; and the duplicated *Lox4* gene was subsequently lost. In the lineage leading to *Cri. mucedo* (Phylactolaemata), the main cluster *Post2* was lost, and the chromosome containing ALG β underwent fission, leaving the main cluster on chr2 and duplicated genes on chr1. In the lineage leading to *M. membranacea* (Gymnolaemata), the duplicated *Lox5* and *Post2* were lost, and the chromosome containing ALG β underwent fission, leaving the main cluster on chr1 and duplicated genes on chr2.

Given the critical dependence of Hox gene regulation on genomic organization, we hypothesized that physical separation might alter gene expression. To investigate this, we analyzed the expression of bryozoan Hox paralogs located on different chromosomes using published RNA-sequencing data (Supplemental Fig. S13; Supplemental Table S14). In *Cri. mucedo*, both the *Hox4b* and *Lox5b* genes separated from the main cluster are expressed more highly than their main cluster counterparts, *Hox4a* and *Lox5a* (paired *t*-tests *p* = 0.0009 and *p* = 0.0001, respectively) (Supplemental Fig. S13). Therefore, the physical separation of Hox cluster genes by genome rearrangements is associated with differences in expression levels.

## Discussion

Our study reveals dramatic rearrangements to bryozoan chromosomes that have extensively fragmented the 24 ALGs inherited by all bilaterians. This presents a stark contrast to reports of ALG conservation from other spiralians (Wang et al. 2017; Martín-Zamora et al. 2023), deuterostomes (Lin et al. 2024), and even non-bilaterians like cnidarians and ctenophores (Simakov et al. 2022; Schultz et al. 2023). We distinguish two distinct periods in which the bryozoan genome was drastically remodelled: the first generating the derived bryozoan ancestral state of just six ALGs from the 24 bilaterian ALGs, largely by chromosome fusion events, and the second independently modifying this state further in different bryozoan classes, including ALG fission events.

Three aspects of bryozoan genomes stand out as particularly remarkable. Firstly, the magnitude of chromosome fusions is notable, with 18 out of 24 (75%) bilaterian ALGs on the same chromosomes. Secondly, the occurrence of these massive fusion events across a phylum is exceptionally rare. Unlike dipterans and clitellate annelids, where such extensive fusion events are specific to certain lineages (Zdobnov et al. 2005; Simakov et al. 2022; Schultz et al. 2024; Lewin et al. 2024a; Vargas-Chávez et al. 2024), our data indicate that these rearrangements are present across the entire extant bryozoan phylum. This suggests that massive rearrangements had already occurred by the time of the last common ancestor of living lineages. However, we note that much of bryozoan diversity is represented only in the fossil record. Six of the seven orders of Stenolaemata are long extinct and less than 40% of bryozoan genera have living representatives (Taylor and Waeschenbach 2015), while approximately 70% of described bryozoan species are fossil (Gordon and Costello 2016). Thirdly, many ALGs in bryozoans have been separated by ALG fissions. Major studies on animal chromosome evolution suggested that a general rule in metazoans is that ALGs are combined but not broken up (Simakov et al. 2022; Wright et al. 2024). In contrast, we identified numerous fissions occurring in the common ancestor of the four gymnolaemate species around 350 million years ago (Orr et al. 2022), with subsequent, lineage-specific fission in *Cry. pallasiana* less than 90 million years ago (Orr et al. 2022). This indicates a prolonged period during which fissions were evolutionarily tolerated, challenging the notion that they occur in brief, time-limited periods and are followed by evolutionary stasis (Simakov et al. 2022).

Why have bryozoans specifically undergone radical ALG fissions while many animal phyla retained the conserved ancestral state? Although our study doesn’t offer a definitive answer, we can apply Mayr’s classical framework (Mayr 1961) to outline the elements of a possible explanation. Firstly, we must identify the mechanism that generated these rearrangements (the proximate cause). Secondly, we must understand why these rearrangements conferred a fitness advantage and were thus favored by selection (the ultimate cause). The high level of rearrangement in bryozoans could be due to either (i) an increase in the rate of rearrangement events (the proximate cause), (ii) an increase in the rate of fixation of rearrangement events (the ultimate cause), or both. Loss of DNA repair machinery (Vargas-Chávez et al. 2024), transposable element invasions (Ahola et al. 2014; Höök et al. 2023), or environmental mutagenic agents (Schultz et al. 2024) have been proposed as potential proximate causes other systems, though additional studies are needed to test whether these were contributors in bryozoans.

The ultimate cause of large-scale genome restructuring events is particularly intriguing from an evolutionary standpoint. One possibility is that rearrangements are selectively neutral or weakly deleterious but are fixed at an increased rate due to genetic drift (Wright 1941; Mackintosh et al. 2023c). Notably, bryozoans are colonial and have an asexual stage in which they reproduce by budding (Schwaha 2020), and genetic drift and mutational load are elevated in asexual populations (Haag and Roze 2007). Alternatively, rearrangements may be adaptive and favored by positive selection, as has been suggested for loss of genome structure during the marine–freshwater transition in clitellate annelids (Sun et al. 2021; Schultz et al. 2024; Vargas-Chávez et al. 2024; Lewin et al. 2024a). The transition to obligate colonialism is a significant evolutionary innovation in bryozoans (Schwaha et al. 2020) that correlates with the timing of genome restructuring, but further work is required to understand whether this, or other bryozoan adaptations such as biomineralization (Taylor et al. 2015), were facilitated by chromosomal rearrangement events.

A third possibility is that increased rate of fixation of rearrangements results from the relaxation in bryozoans of strong evolutionary pressure that has been maintained in other lineages. The role of conserved macrosyntenic gene linkages in genome functioning remains largely unknown, but their high-fidelity maintenance in diverse lineages for over half a billion years (Simakov et al. 2022) implies at least some importance. One idea is that this is due to long-range interactions between distant elements on the same chromosome, which means that gene linkages are significant to gene expression (Tolhuis et al. 2002; Miele and Dekker 2008; Dean 2011). A change in the relative importance of mechanisms of gene regulation may therefore reduce the emphasis on long-range interactions and render conserved genome organization obsolete. Our work highlights the need for studies relating gene regulation to genome organization, which may be key to understanding the ultimate causes of the evolution of highly derived genomic architectures.

Having elucidated the organization of bryozoan genomes, we questioned whether macrosynteny could resolve their phylogenetic position. Substantial evidence has been presented supporting both their association with Phoronida and Brachiopoda (Lophophorata) (Nesnidal et al. 2013; Luo et al. 2017; Marlétaz et al. 2019; Laumer et al. 2019) and with Entoprocta and Cycliophora (Polyzoa) (Hejnol et al. 2009; Kocot et al. 2017; Khalturin et al. 2022). Our study identifies nine fusion-with-mixing events shared between bryozoans and brachiopods, of which five are confirmed as derived characters, supporting the Lophophorata topology. Fusion-with-mixing events are essentially irreversible in evolution, but convergent evolution cannot be ruled out (Steenwyk and King 2024). If these rare genomic changes are specific to Lophotrochozoans (excluding Entoprocta and Cycliophora), then the presence of at least nine shared derived fusion events in brachiopods and bryozoans would almost rule out convergence, becoming a conclusive piece of evidence in this ongoing debate. The future availability of chromosome-scale genomes from these groups is expected to provide clarity. Looking further, evidence from macrosynteny is likely to become increasingly prominent in discussions concerning phylogenetically challenging clades.

Overall, though macrosyntenic gene linkages are conserved across diverse bilaterian lineages, members of the phylum Bryozoa possess highly derived genomes with scrambling of these gene sets. Divergence has continued within the phylum, leading to distinct organizational structures in two bryozoan classes, and ALG fission, previously underappreciated in its importance, has occurred with high frequency. The addition of information on the third bryozoan class (the Stenolaemata, represented by present-day cyclostomes) when chromosome-level genomes become available will add considerably to this picture. Furthermore, although we present comparative genomics data on Hox gene duplications, additional studies on the spatial expression patterns of bryozoan Hox genes are necessary to examine how genome rearrangements have affected their functions (Fuchs et al. 2011). We expect the bryozoan genomes annotated in this study to be crucial for the pursuit of such future research avenues.

## Methods

### Assembly acquisition and gene prediction

Chromosome-scale assemblies of *B. stolonifera* (GenBank accession no. GCA_935421135) (Wood et al. 2023), *Cry. pallasiana* (GCA_945261195) (Bishop et al. 2023b), *M. membranacea* (GCA_914767715) (Bishop et al. 2023a), and *W. subatra* (GCA_963576615) (Bishop et al. 2024) were sequenced as part of the Darwin Tree of Life project (The Darwin Tree of Life Project Consortium et al. 2022) and retrieved from the NCBI Datasets resource. A chromosome-scale assembly of *Cri. mucedo* (Hoencamp et al. 2021) was obtained from the NCBI Gene Expression Omnibus (GSE169088). None of the bryozoan assemblies used have publicly available gene models, so gene prediction was performed in-house. Repetitive elements were first annotated with RepeatModeler (v2.0.4) (Flynn et al. 2020) and masked using RepeatMasker (v4.1.5) in sensitive mode (Smit et al. 2015). Gene prediction and annotation were performed using the BRAKER3 pipeline (v3.0.3), incorporating hints from RNA sequencing (RNA-seq) data as aligned reads in bam format (Stanke et al. 2006, 2008; Li et al. 2009; Barnett et al. 2011; Lomsadze et al. 2014; Buchfink et al. 2015; Hoff et al. 2016, 2019; Gabriel et al. 2024).

BRAKER3 uses RNA-seq and genome data to train the gene prediction tools AUGUSTUS and GeneMark, producing highly supported gene transcripts. Raw RNA-seq reads for gene prediction (Supplemental Table S2) were sourced from the NCBI Sequence Read Archive (SRA) using the SRA Toolkit (v3.0.0) (Leinonen et al. 2011) with GNU parallel (Tange 2021). Transcriptomic data for *B. stolonifera*, *Cry. pallasiana*, *M. membranacea*, and *W. subatra* were generated from adult colonies collected by the Marine Biological Association, Plymouth, UK, in association with the Darwin Tree of Life project. For these four species, RNA extraction was performed on the same colony used for DNA sequencing to generate the reference assembly. For *B. stolonifera* and *M. membranacea,* this dataset was supplemented with independent datasets from developmental stages (Supplemental Table S2). RNA-seq data for *Cri. mucedo* was derived from mature adult colonies from a population different to that from which the genome assembly was produced (Supplemental Table S2). Quality control for RNA-seq data was performed with FastQC (Andrews and Others 2010). Reads were trimmed with Trimmomatic (v0.39) (Bolger et al. 2014) and aligned to the genome with STAR (v2.7.10b) (Dobin et al. 2013). BUSCO (v5.4.7) (Simão et al. 2015; Manni et al. 2021) was used to evaluate the quality of gene prediction. Hox genes were identified with reciprocal tBLASTn searches (Camacho et al. 2009), and orthology was confirmed using gene trees built in IQ-TREE (v2.2.2.3) (Minh et al. 2020). Hox annotations are consistent with previous work, where available (Saadi et al. 2023).

### Phylogenetics

Single-copy orthologs were identified using OrthoFinder (v2.5.4) (Emms and Kelly 2019). Alignments for each set of orthologs were created with MAFFT (v7.520) (Katoh et al. 2002; Katoh and Standley 2013), trimmed with ClipKIT (v1.4.1) (Steenwyk et al. 2020), and concatenated with PhyKIT (v1.11.7) (Steenwyk et al. 2021). IQ-TREE (v2.2.2.3) (Minh et al. 2020) was used to construct maximum likelihood trees with automated substitution model selection using ModelFinder (Kalyaanamoorthy et al. 2017) and 1,000 UFBoot2 ultra-fast bootstrap replicates (Hoang et al. 2018). Trees were rendered with iTOL (Letunic and Bork 2021). A tree of Hox protein sequences was generated using the same IQ-TREE method.

### Macrosynteny analysis

We developed SyntenyFinder, a pipeline comprising a collection of Python and R scripts designed to facilitate macrosynteny analysis using chromosome-level genome assemblies. SyntenyFinder is a straightforward pipeline with two main functions: first, it uses OrthoFinder (v2.5.4) (Emms and Kelly 2019) to identify single-copy orthologs in a dataset of protein sequence files. Second, it utilizes a custom R script to generate chromosome idiograms using RIdeogram (v0.2.2) (Hao et al. 2020) and to create dot plots showing the positions of orthologous genes between two genomes. SyntenyFinder is made available on Github (https://github.com/symgenoevolab/SyntenyFinder) and can be run in two modes: either automatically downloading and running on annotated genomes from NCBI by their accessions, or on local files parsed directly to the program. It requires three types of file as an input: (1) genome sequence FASTA file; (2) gene positions in the format of a GTF file; and (3) a protein sequence FASTA file.

For the purposes of this paper, two runs were performed: first with only the five bryozoan species, and second adding *B. floridae* (Chordata), *P. maximus* (Mollusca), *L. longissimus* (Nemertea) and *L. anatina* (Brachiopoda). In both cases, only orthologs which were single-copy in all study species were used for downstream analysis. No gene loss or duplication was permitted. In the second case, to track the genomic position of bilaterian ALGs, we used an input dataset of the genes used to define the 24 bilaterian ALGs (Simakov et al. 2022) and used OrthoFinder to find orthologs of these genes in bryozoan genomes. The restriction to genes with known ALG relationships reduces the noise within the dataset. Macrosynteny plots for this manuscript were created in R (v4.3.0) (R Core Team 2023). The same set of genes is used for each comparison in both idiograms and dot plots of macrosynteny. In idiograms, links between chromosomes with fewer than five shared genes were removed for visual clarity. These represent single-gene or few-gene translocations. All genes, including translocated genes, are shown in dot plots. For plotting purposes and to ensure alignment with orthologous chromosomes, the orientations of chr1, chr2, chr4, chr6, chr7, chr10, and chr11 in *W. subatra*, and chr2, chr3, chr7, chr9, chr10, chr11, chr12 in *Cry. pallasiana* were reversed from their orientations as presented in published datasets.

### Examination of large-scale duplications

Two complementary methods were used to determine whether segmental duplication or whole genome duplication (WGD) events have occurred in the bryozoan lineage. In each case, we use the annelid *M. vulgaris* as a positive control for a species with a known WGD (Jin et al. 2020; Lewin et al. 2024a; Vargas-Chávez et al. 2024). First, synonymous substitutions per site (*K_s_*) distributions were calculated with wgd (v.2.0.26) (Chen et al. 2024). Briefly, paralogs were identified with ‘wgd dmd’ using Diamond (v2.0.14.152) (Buchfink et al. 2015, 2021) and then ‘wgd ksd’ was run to calculate the *K_s_* distribution using PAML (v4.9) (Yang 2007). The *K_s_* between two paralogs is an estimate of their divergence time. The null expectation with no WGD is that the number of paralogs decreases exponentially as *K_s_* increases. When a WGD occurs, an excess of duplicates are created simultaneously, creating a peak in the distribution (Tiley et al. 2018). Paralogs situated in collinear blocks in the genome, so-called ‘anchor duplicates’, are more likely to be produced by WGD than dispersed paralogs and are therefore more reliable for WGD detection (Chen et al. 2024). Anchor duplicates were identified using ‘wgd syn’ to implement the collinearity search algorithm of i-ADHoRe (v3.0.01) (Proost et al. 2011). *K_s_* distributions using only anchor duplicates were then plotted with ‘wgd syn’ (Chen et al. 2024).

Second, MCScanX (Wang et al. 2012) was used to identify collinear blocks of duplicated genes within bryozoan genomes and to classify all duplicates within each genome as dispersed, proximal, tandem or WGD/segmental. The presence of large homologous regions within a genome containing collinear blocks of duplicated genes is suggestive of WGD. The MCScanX output was plotted as a Circos plot (Krzywinski et al. 2009) using shinyCircos (v2.0) (Wang et al. 2023).

### Validation of chromosome fission

Given the absence of a high-quality Hi-C contact map for *Cry. pallasiana* (Bishop et al. 2023b), the fission event in this species remained uncertain. To validate the fission event that led to the formation of chromosomes CPA10 and CPA12, as shown in Fig. 2, we examined chromatin interactions by mapping Hi-C data to the chromosome-level assembly. Hi-C reads for *Cry. pallasiana* were downloaded from the NCBI SRA (accession ERR9866429) using the SRA Toolkit. The SRA file was fetched with prefetch and converted to FASTQ format using fasterq-dump. The Hi-C reads were then aligned to the genome assembly (accession GCA_945261195.1) using BWA (v0.7.17-r1198) (Li 2013), following the Arima Hi-C mapping pipeline (A160156 v03). A Hi-C contact map was generated using YaHS (v1.2a.1) (Zhou et al. 2023) and JuicerTools (v1.9.9), and visualized with JuiceBox (v1.11.08) (Durand et al. 2016).

### Quantification of microsynteny mixing score

To assess the amount of intra-chromosomal gene shuffling, we first calculated Spearman’s rank correlation coefficient (*ρ*). This coefficient quantifies the strength and direction of association between the ranked order of orthologous genes along orthologous chromosome pairs. The Spearman’s coefficient is given by the following equation:

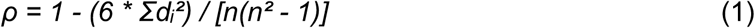

where *ρ* is the coefficient, *dᵢ* represents the difference in ranks between corresponding values of the two variables, and *n* is the total number of observations. Subsequently, we defined the synteny mixing score, an indicator of chromosomal rearrangement, using the equation:

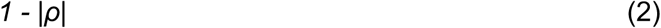

where *|ρ|* denotes the absolute value of the Spearman’s coefficient. This metric allows the synteny mixing score to vary from 0, indicative of no intra-chromosomal rearrangement and thus a high degree of conserved microsynteny, to 1, representative of maximum rearrangement, signifying considerable disruption in the original gene order.

### Estimation of species divergence time

A phylogenetic tree, constructed above from single-copy orthologs extracted from five bryozoan genomes and a mollusc outgroup (*P. maximus*), was used to estimate divergence times. We calibrated this tree with contemporary molecular and fossil data (Orr et al. 2022), focusing on the divergence between *P. maximus* and *Cri. mucedo* during the Cambrian Period (541 to 488 million years ago) (Erwin et al. 2011), employing the ape package in R (Paradis and Schliep 2018).

### Quantification of gene expression

Gene expression was quantified in a published *Cri. mucedo* (PRJNA594616) RNA-seq datasets. Raw reads were obtained from the NCBI SRA using the SRA toolkit (Leinonen et al. 2011) and GNU parallel (Tange 2021). Hox gene expression was quantified with Salmon (v1.10.2) (Patro et al. 2017). *Hox4a*, *Hox4b*, *Lox5a*, and *Lox5b* gene expression levels were compared using *t*-tests with a Benjamini-Hochberg correction for multiple testing.

### Data access

Gene models for all five bryozoan species have been deposited in Dryad (https://doi.org/10.5061/dryad.76hdr7t3f). Custom Python and R scripts used for macrosynteny analysis, calculating mixing scores, and estimating divergence times are available in our GitHub repository (https://github.com/symgenoevolab/SyntenyFinder) and as Supplemental Code.

## Acknowledgments

This work was funded by a Royal Society Newton International Fellowship (NIF\R1\201315) and an Academia Sinica Career Development Award (AS-CDA-112-L06) to Y.-J.L. We acknowledge the efforts in sample collection by the Marine Biological Association (MBA)– Darwin Tree of Life collecting team: Patrick Adkins, Robert Mrowicki, Joanna Harley, and Helen Jenkins, as well as Christine Wood of the MBA. We thank the members of the Symbiosis Genomics & Evolution Lab for their assistance and support.

## Author contributions

T.D.L. and Y.-J.L. designed research. T.D.L., I.J.-Y.L., M.-E.C., and Y.-J.L. performed research. T.D.L., I.J.-Y.L., J.D.D.B., and Y.-J.L. contributed new reagents/analytic tools. T.D.L. and Y.-J.L. analyzed data. T.D.L., P.W.H.H., and Y.-J.L. wrote the paper. All authors read and approved the final manuscript.

