## Supplemental Material for "Fusion, fission, and scrambling of the bilaterian genome in Bryozoa"

**This PDF file includes:**

Supplemental Figs. S1 to S13
Supplemental Tables S1 to S14
Legends for Supplemental Data S1 to S2
Supplemental References

**Other supporting materials for this manuscript include the following:**

Supplemental Data S1 to S2

**Table of contents**

***Supplemental Figures*** ***3***

*Supplemental Fig. S1* 3
*Supplemental Fig. S2* 4
*Supplemental Fig. S3* 5
*Supplemental Fig. S4* 6
*Supplemental Fig. S5* 7
*Supplemental Fig. S6* 8
*Supplemental Fig. S7* 10
*Supplemental Fig. S8* 11
*Supplemental Fig. S9* 12
*Supplemental Fig. S10* 13
*Supplemental Fig. S11* 14
*Supplemental Fig. S12* 15
*Supplemental Fig. S13* 16

***Supplemental Tables*** ***17***

*Supplemental Table S1* 17
*Supplemental Table S2* 18
*Supplemental Table S3* 19
*Supplemental Table S4* 20
*Supplemental Table S5* 22
*Supplemental Table S6* 23
*Supplemental Table S7* 24
*Supplemental Table S8* 26
*Supplemental Table S9* 28
*Supplemental Table S10* 30
*Supplemental Table S11* 31
*Supplemental Table S12* 32
*Supplemental Table S13* 33
*Supplemental Table S14* 34

***Legends for Supplemental Datasets*** ***35***

***Supplemental References*** ***35***

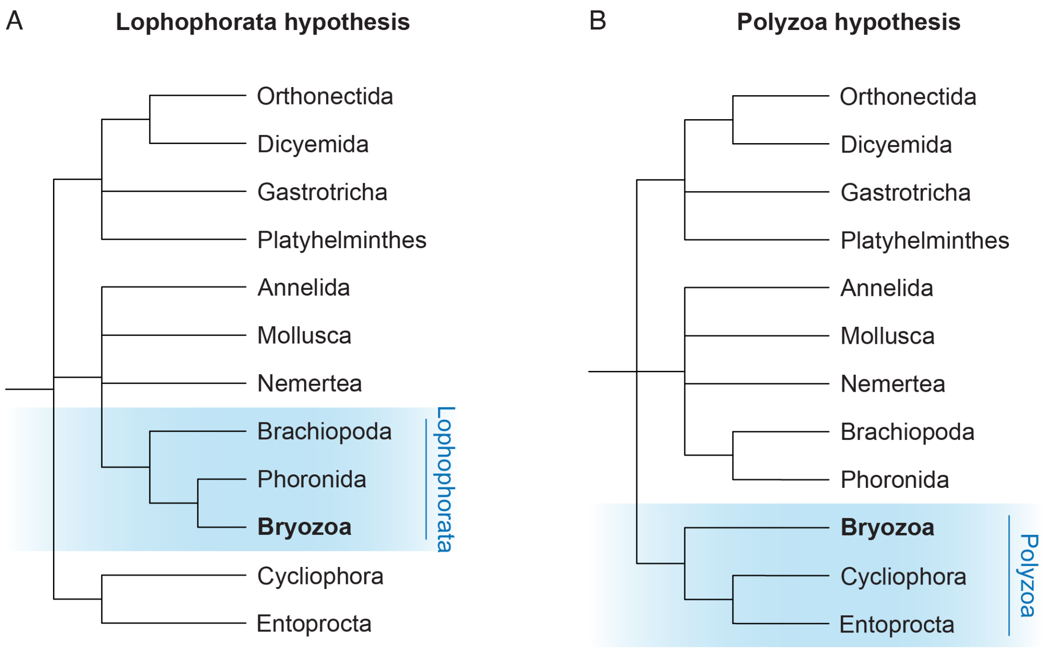


**Supplemental Fig. S1. Lophophorata and Polyzoa hypotheses for the phylogenetic position of Bryozoa.** (*A*) Phylogeny of spiralian phyla under Lophophorata hypothesis. (*B*) Phylogeny of spiralian phyla under Polyzoa hypothesis. Consensus phylogenies are from Liao et al. (2023).

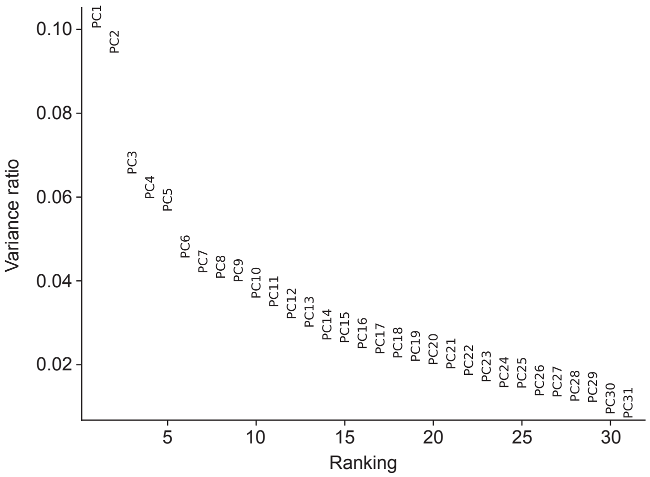


**Supplemental Fig. S2. Variance ratios for the top 30 principal components for gene family analysis.** Main figure for principal component analysis is Main Text Fig.1E.

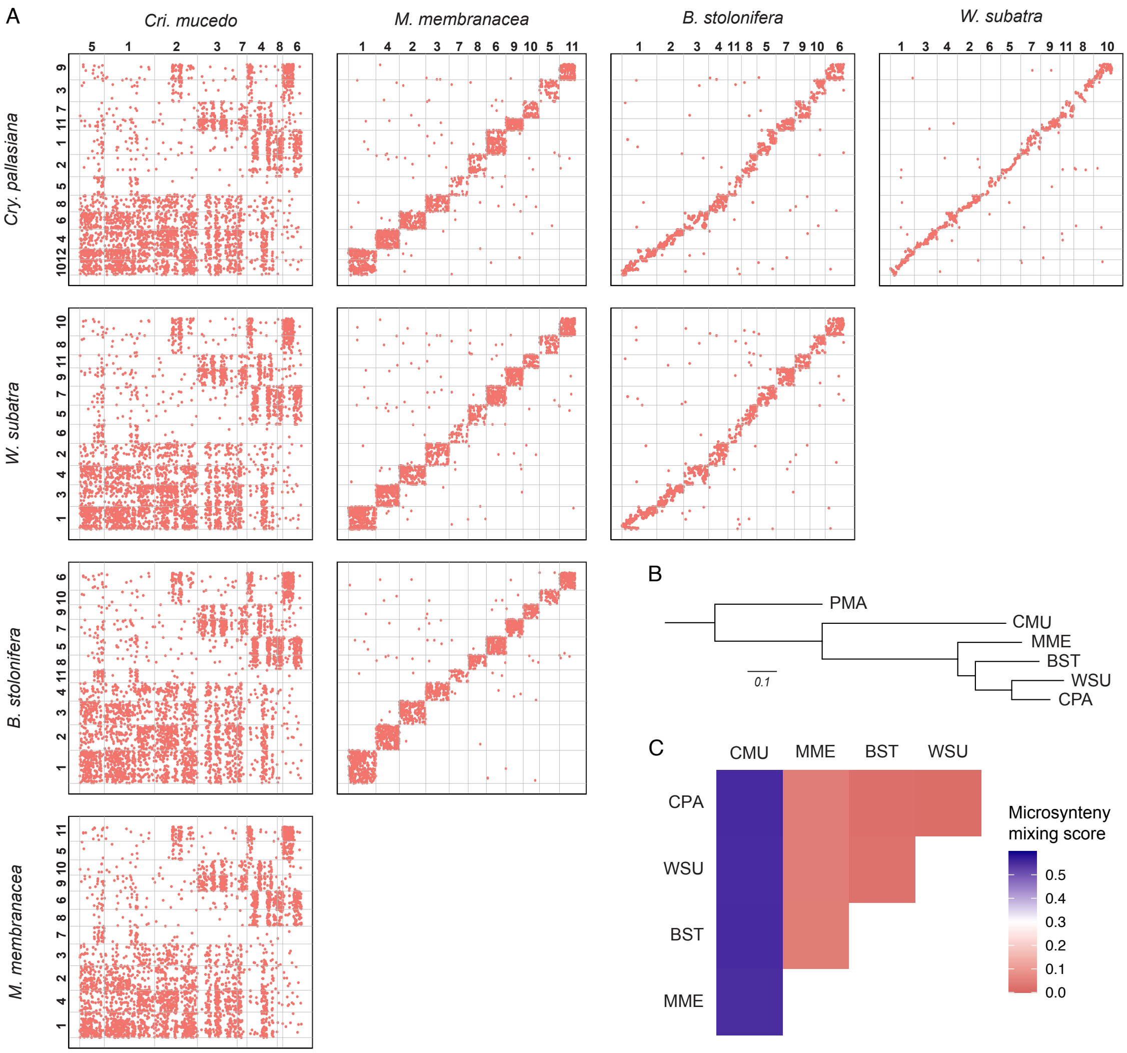


**Supplemental Fig. S3. Chromosome evolution within bryozoans revealed by Oxford dot plots.** (*A*) Oxford dot plots show pairwise comparisons of the position of orthologs in two genomes. Red points represent orthologous genes, axes represent entire genome length. Pale gray bars mark chromosome boundaries. Bold numbers show chromosome number. A straight line from bottom left to top right would represent complete conservation of macrosynteny and gene order (*W. subatra* vs. *Cry. pallasiana* is the closest). The widening of the line into square blocks of genes represents the shuffling of orthologous genes within orthologous chromosomes (e.g. *M. membranacea* vs. *Cry. pallasiana*). Blocks covering one chromosome in one species but multiple chromosomes in another species represent fusion or fission events (e.g. *Cri. mucedo* vs. *Cry. pallasiana*). *Cry. pallasiana*, *W. subatra*, *B. stolonifera* and *M. membrancea* have highly conserved genome structure. The *Cri. mucedo* genome is highly rearranged in comparison. (*B*) Phylogeny of bryozoan study species reproduced from Main Text Figure 1C. (*C*) Quantification of microsynteny (shuffling of genes within chromosomes) between species with our 'microsynteny mixing score'. More closely related species have a lower microsynteny mixing score, indicating lower rates of shuffling and more highly conserved gene order. Scores are listed as Table S9.

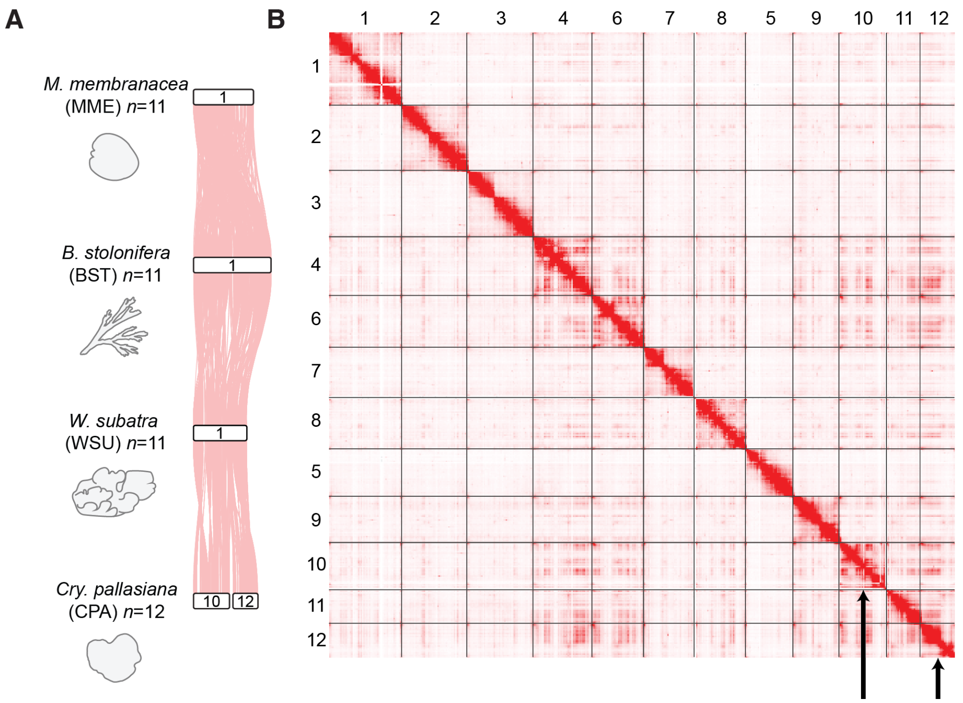


**Supplemental Fig. S4. Macrosynteny analysis together with Hi-C contact map for *Cryptosula pallasiana* reveals chromosome fission.** (*A*) Synteny analysis across bryozoan genomes showed lineage-specific chromosome fission. (*B*) *Cry. pallasiana* chromosomes 10 and 12 (arrows) are clearly separated on the contact map, demonstrating that these chromosomes are the products of a genuine chromosome fission and not a mis-assembly. Darker colors indicate a higher frequency of contacts.

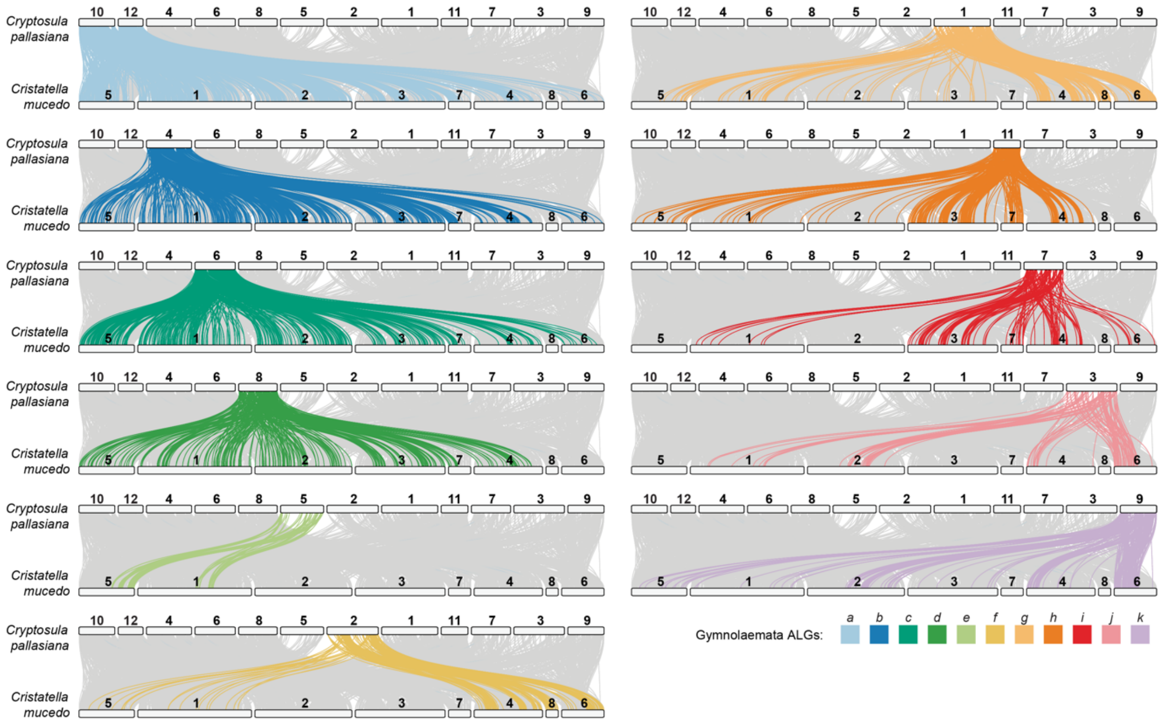


**Supplemental Fig. S5. Relationships between *Cry. pallasiana* and *Cri. mucedo* chromosomes.** Chromosome-scale gene linkage between *Cry. pallasiana* (representative of Gymnolaemata) and *Cri. mucedo* (Phylactolaemata) represented by ribbon plots. Horizontal bars represent chromosomes. Vertical lines connect the genomic position of orthologous genes in each genome. Each plot highlights genes from one Gymolaemata ancestral linkage group (ALG). In Gymnolaemata species, 11 ALGs are highly conserved, though the chromosome containing ALG *a* underwent a fission event to form chr10 and chr12 in *Cry. pallasiana*. These ALGs are not conserved in the Phylactolaemata species *Cri. mucedo*, and are distributed across the genome.

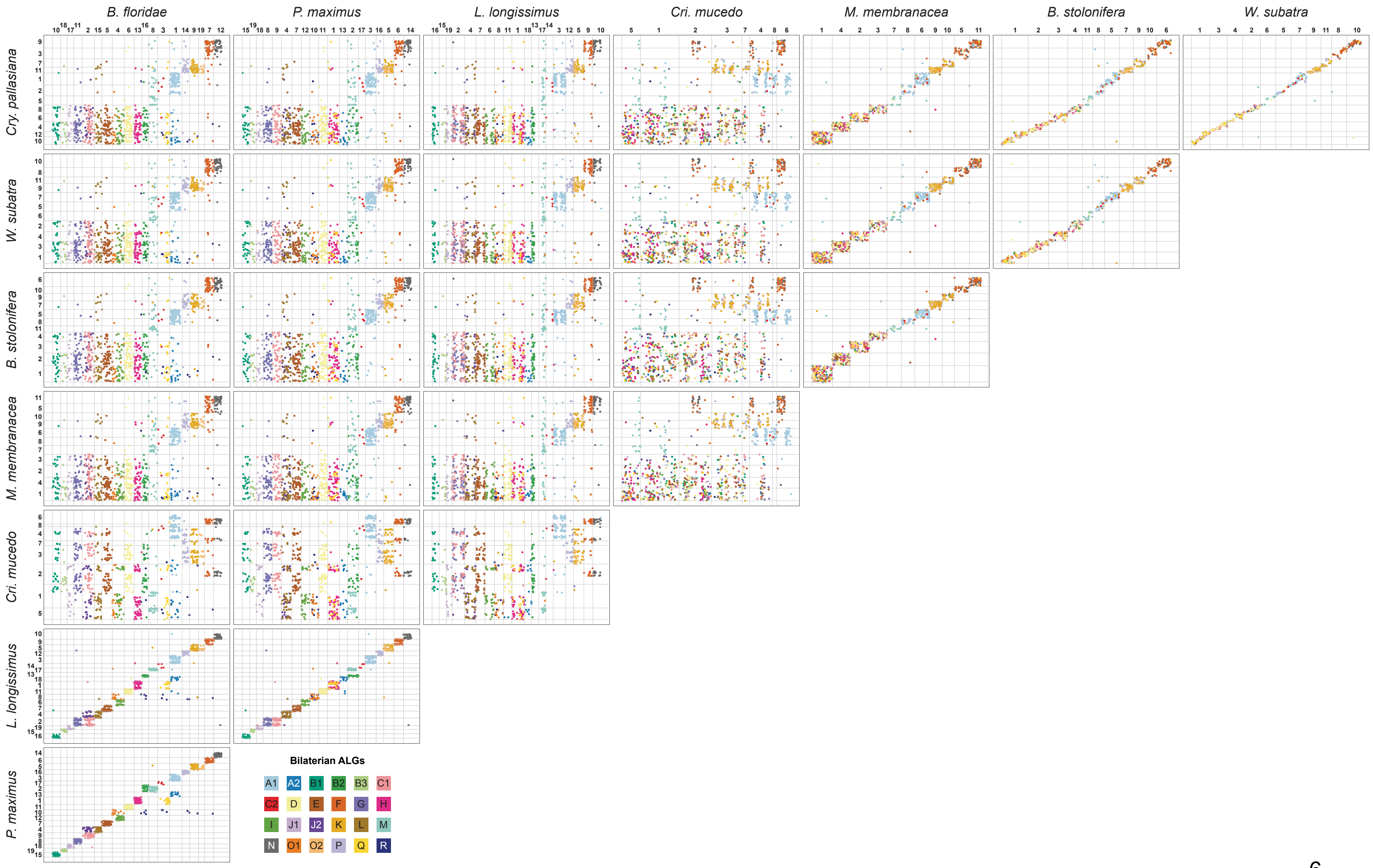
B. floridae
P. maximus
L. longissimus
Cri. mucedo
M. membranacea
B. stolonifera
W. subatra
Cry. pallasiana
Bilaterian ALGs
A1
A2
B1
B2
B3
C1
C2
D
E
F
G
H
I
J1
J2
K
L
M
N
O1
O2
P
Q
R

**Supplemental Fig. S6. Chromosome relationships between bryozoans, spiralians and other bilaterians revealed by Oxford dot plots.** Oxford dot plots show chromosome-scale gene linkage for species used in this work. Each point represents a gene. Genes are colored by their bilaterian ALG. Gray lines mark chromosome boundaries. Macrosynteny is highly conserved between *B. floridae*, *P. maximus* and *L. longissimus*, reflecting the bilaterian ancestral state. Macrosynteny is also highly conserved within Gymnolaemata bryozoans, but this state represents a substantial rearrangement from the bilaterian ancestral state.

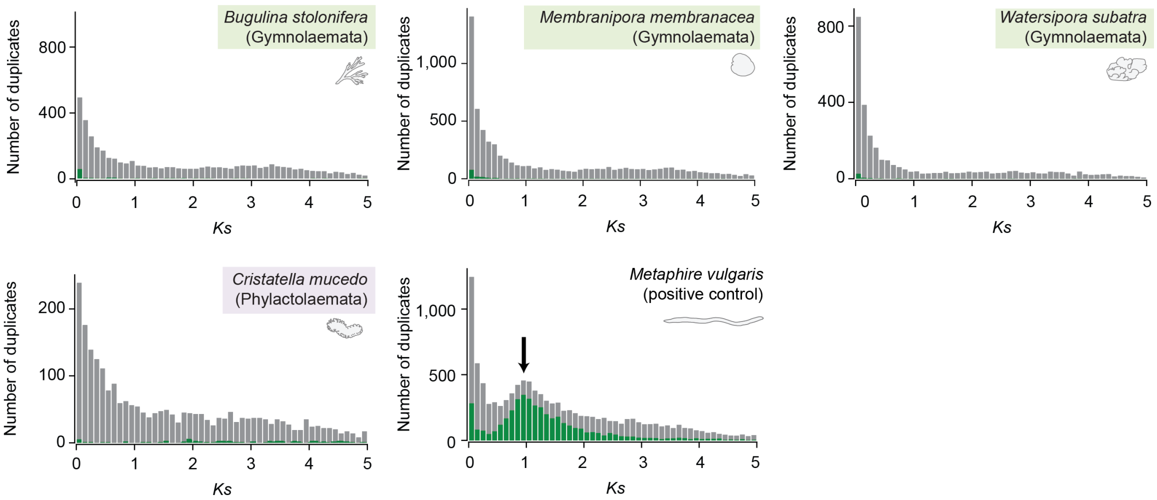


**Supplemental Fig. S7. Assessment of whole genome duplication (WGD) in bryozoan genomes using synonymous substitution rate (*K<sub></sub>S*).** Gray bars show histograms for all duplicated genes, while green bars show only anchor duplicates: these are duplicated genes found in duplicated collinear blocks of genes in the genome. Genomes with no WGD are expected to show exponential decay of duplicate numbers as $K_S$ increases. *M. vulgaris* is a positive control with a known WGD (Jin et al. 2020): the plot for this species is interrupted by a large normally distributed peak at $K_S = 1$, indicating recent WGD (arrow). The plots of the bryozoans have no similar large peak at a low $K_S$, suggesting no recent WGD. There possible presence of a very shallow, very broad peak at a higher $K_S$ in gymnolaemate bryozoans means that WGD in this lineage cannot conclusively be ruled out. The phylactolaemate bryozoan *Cri. mucedo* has several very small peaks of anchor gene duplicates (green), possibly representing segmental duplication events.

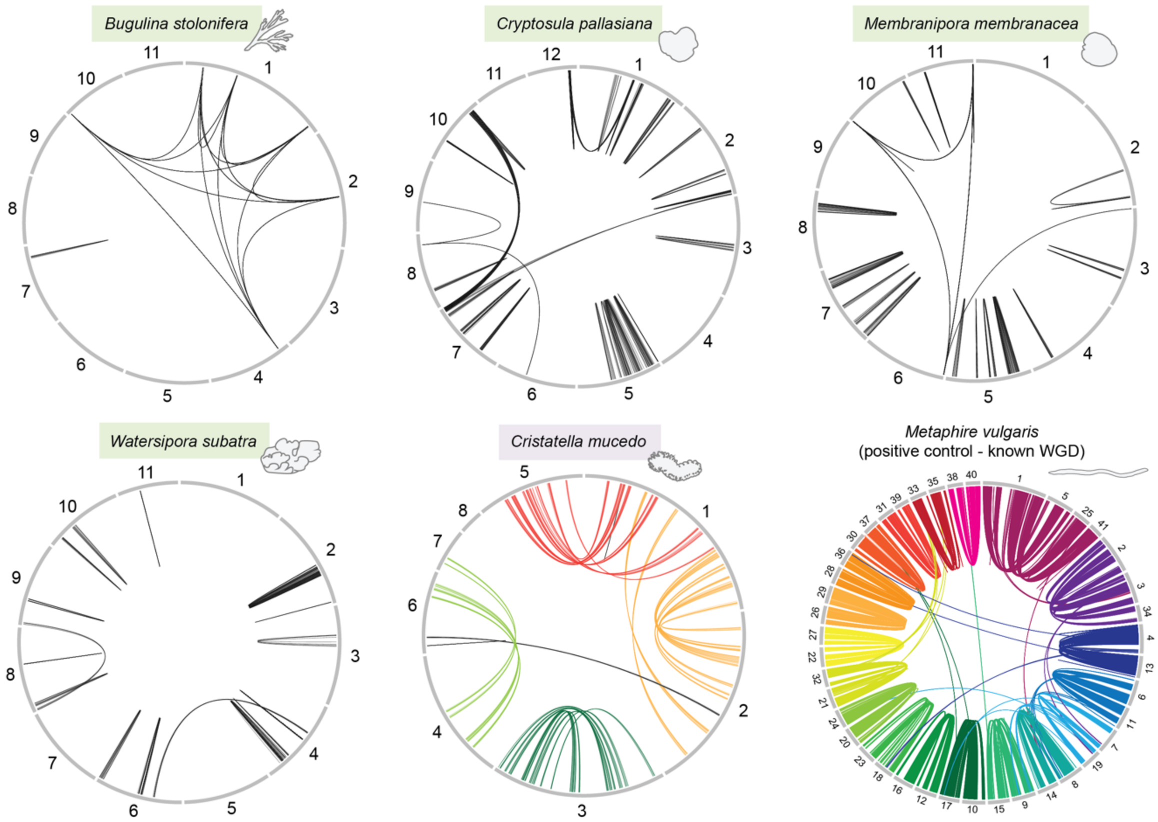


**Supplemental Fig. S8. Duplicated collinear gene blocks in bryozoan genomes.** Duplicated collinear blocks indicate that chromosomes likely emerged through WGD or large segmental duplications. *M. vulgaris* is a positive control with a known WGD: the plot for this species shows clear pairs of chromosomes with conserved collinear gene blocks; these emerged from WGD. The plots for gymnolaemate bryozoans show no such blocks, suggesting no WGD. The plot for the phylactolaemate *Cri mucedo* has several sections of homology (chromosome 1 with 2; 1 with 5; 4 with 6), suggesting possible segmental duplication or WGD events in this lineage.

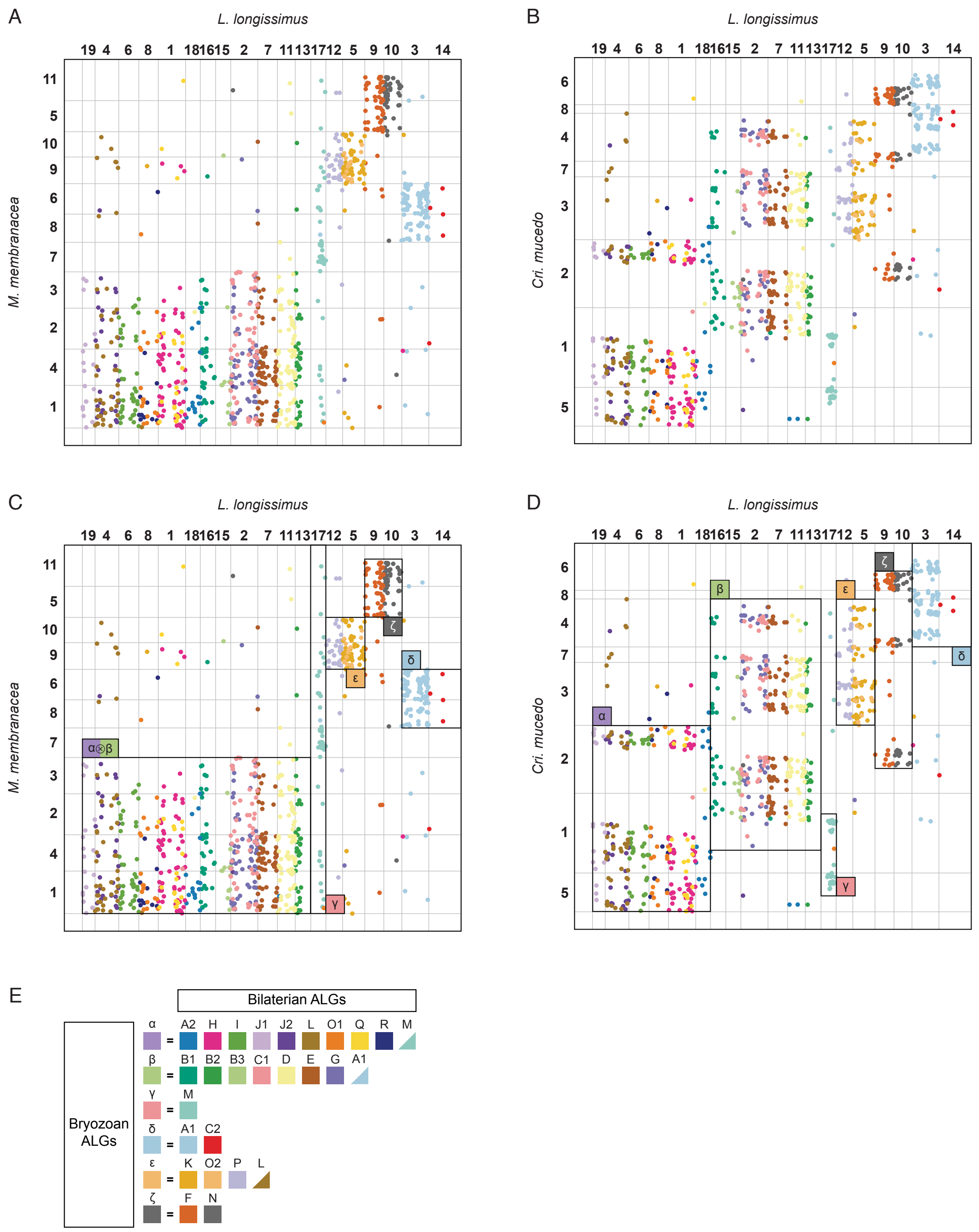


**Supplemental Fig. S9. Oxford dot plots reveal bryozoan ALGs.** Oxford dot plots show chromosome-scale gene linkage of *L. longissimus*, which closely resembles the bilaterian ancestral state with (*A*) *M. membranacea* and (*B*) *Cri. mucedo*. Each point represents a gene. Genes are colored by their bilaterian ALG. Gray lines mark chromosome boundaries. Six bryozoan ALGs can be identified by sets of genes grouped together. In parts (*C*) and (*D*), the six bryozoan ALGs are highlighted with black boxes. (*E*) Composition of bryozoan ALGs by bilaterian ALGs reproduced from Main Text Figure 4B.

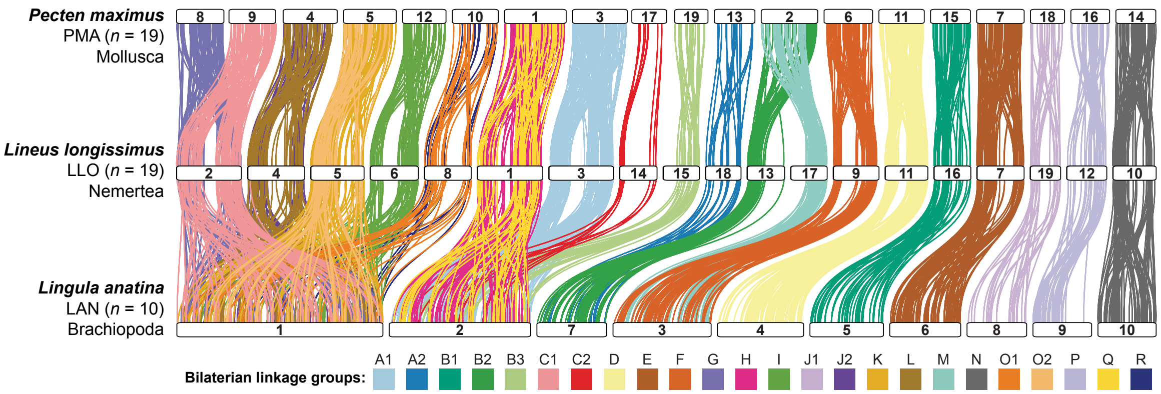


**Supplemental Fig. S10. Chromosome evolution in *L. anatina*.** Chromosome relationships between *P. maximus*, *L. longissimus* and *L. anatina* revealed by ribbon plots. Horizontal white bars represent chromosomes. Vertical colored lines connect orthologous genes on chromosomes. Four *L. anatina* chromosomes (chr1, chr2, chr3, and chr7) are the products of fusion-with-mixing events. Chr1 contains nine bilaterian ALGs (C1⊗G⊗I⊗J2⊗K⊗L⊗O1⊗O2⊗R); chr2 contains five ALGs (A1⊗B3⊗C2⊗H⊗Q); chr3 and chr7 contain two ALGs each (F⊗M and A2⊗B2, respectively). No fission events can be detected in the *L. anatina* assembly.

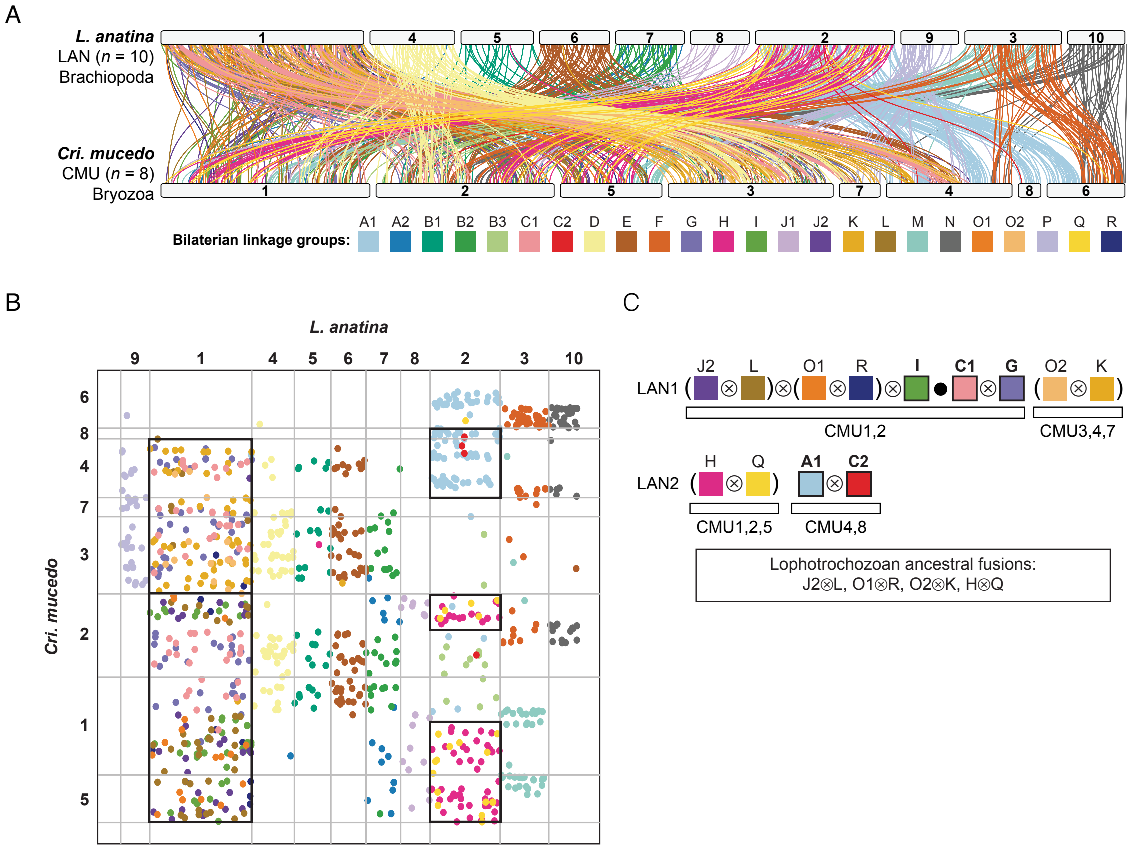


**Supplemental Fig. S11. Shared fusion events between *L. anatina* and *Cri. mucedo*.** (*A*) Chromosome-scale gene linkage between the brachiopod *L. anatina* and the bryozoan *Cri. mucedo*. Horizontal bars represent chromosomes. Vertical lines connect the genomic position of orthologous genes in each genome. Lines are colored by bilaterian ALGs. (*B*) Oxford dot plot revealing chromosome-scale gene linkage between *L. anatina* and *Cri. mucedo*. Black boxes highlight shared fusion-with-mixing events. (*C*) Nine fusion events are shared between *L. anatina* and *C. mucedo*. Brackets surround the four events which are shared by several spiralian phyla and whose timing is uncertain. These events may therefore not be useful phylogenetic markers for the position of bryozoans. The remaining five events are derived fusions shared by bryozoans and brachiopods. In *Cri mucedo*, the C1⊗G group has fused but not mixed (●) with the (J2⊗L)⊗(O1⊗R)⊗I group.

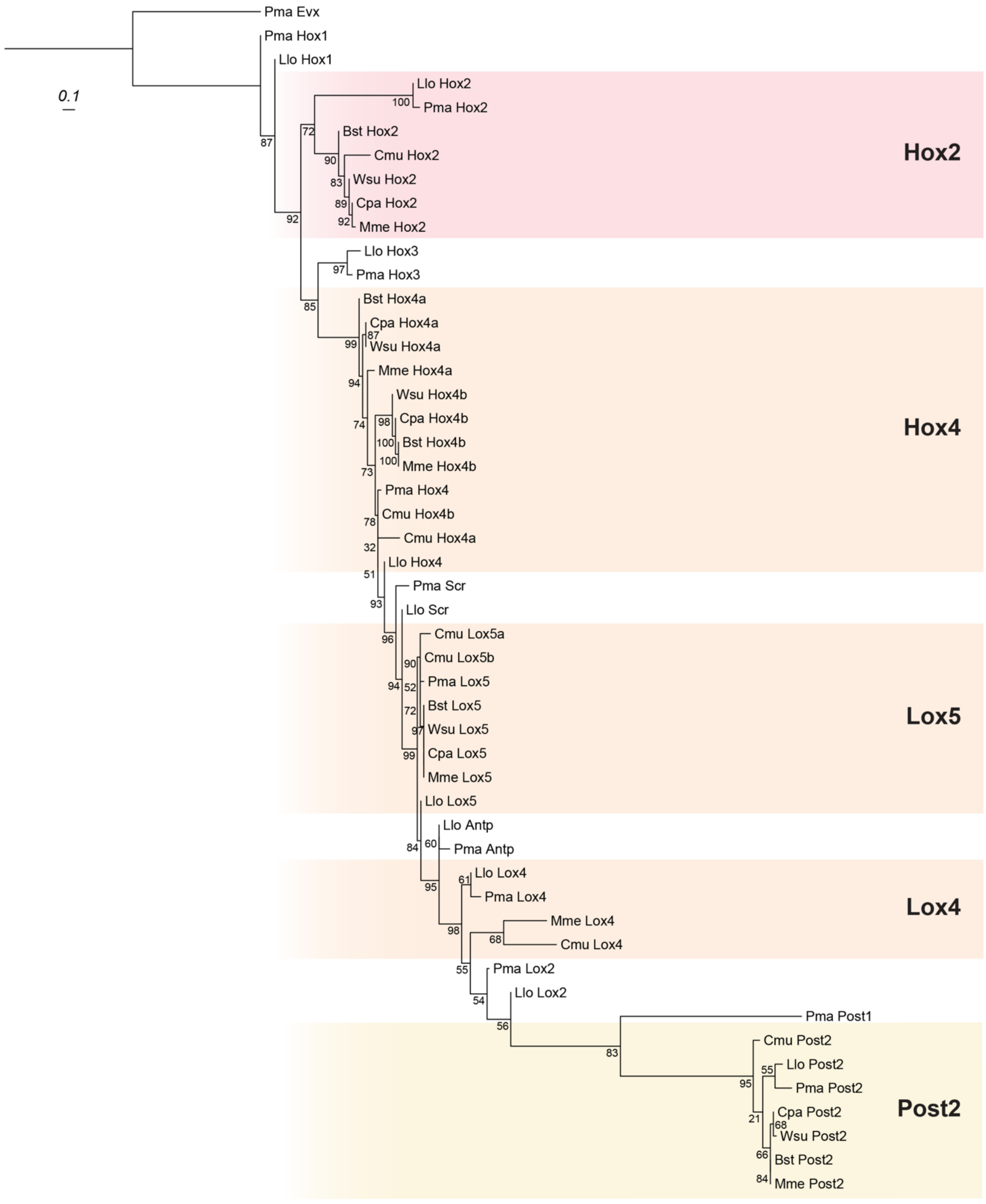


**Supplemental Fig. S12. Tree of bryozoan Hox sequences.** Tree of bryozoan Hox protein sequences used in this analysis. Tree is made using the maximum likelihood method (LG+G4 model) with 1000 bootstrap replicates in IQ-TREE (Minh et al. 2020). *Pecten maximus* Evx is the outgroup.

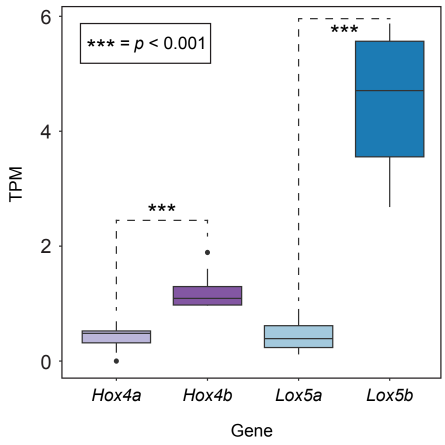


**Supplemental Fig. S13. Expression of *Hox4a* versus *Hox4b* and *Lox5a* versus *Lox5b* in *Cri. mucedo* adults.** Abbreviations: TPM, transcripts per million.

**Supplemental Table S1.** Assembly statistics for bryozoan genomes.

| **Assembly** | ***Cristatella mucedo*** | ***Membranipora membranacea*** | ***Bugulina stolonifera*** | ***Watersipora subatra*** | ***Cryptosula pallasiana*** |
|---|---|---|---|---|---|
| Reference | Hoencamp et al. 2021 | Bishop et al. 2023a | Wood et al. 2023 | Bishop et al. 2024 | Bishop et al. 2023b |
| Assembly level | Chromosome | Chromosome | Chromosome | Chromosome | Chromosome |
| Sequencing technology | PacBio, Illumina, Hi-C | PacBio, 10X Genomics Chromium, Arima Hi-C | PacBio, 10X Genomics Chromium, Arima2 Hi-C | PacBio, 10X Genomics Chromium, Arima Hi-C | PacBio, 10X Genomics Chromium, Arima2 Hi-C |
| Genome size (Mb) | 556 | 339 | 235 | 784 | 606 |
| Chromosome number | 8 | 11 | 11 | 11 | 12 |
| Sequencing coverage | ? | 66X | 52X | 39X | 35X |
| Number of scaffolds | 753 | 21 | 30 | 240 | 847 |
| Scaffold N50 (Mb) | 97 | 30 | 20.3 | 66.5 | 46.3 |
| Contig N50 (Mb) | 0.4 | 7 | 12.6 | 4.5 | 1.8 |
| GC content (%) | 47.0 | 38.0 | 37.0 | 35.5 | 37.2 |
| Repeats (%) | 47.5 | 35.5 | 32.2 | 53.2 | 57.0 |
| Number of genes | 29904 | 25909 | 20502 | 25115 | 30048 |
| Number of transcripts | 32678 | 31934 | 25449 | 21344 | 35283 |
| Mean gene length (bp) | 10823 | 6430 | 6766 | 14392 | 7863 |
| Mean number of introns per gene | 6.0 | 8.6 | 9.1 | 7.9 | 6.9 |
| Mean intron size (bp) | 1704 | 683 | 658 | 1847 | 1005 |
| Genome BUSCO metazoa *odb10* (n = 954) | C:88.6% [S:87.4%, D:1.2%] F:6.2% M:5.2% | C:82.7% [S:81.0%, D:1.7%] F:8.0% M:9.3% | C:84.5% [S:83.8%, D:0.7%] F:6.5% M:9.0% | C:83.4% [S:81.7%, D:1.7%] F:8.6% M:8.0% | C:83.6% [S:82.4%, D:1.2%] F:8.0% M:8.4%, |
| Transcriptome BUSCO metazoa *odb10* (n = 954) | C:86.7% [S:85.3%, D:1.4%] F:6.8% M:6.5% | C:94.9% [S:91.9%, D:3.0%] F:2.6% M:2.5% | C:96.6% [S:95.4%, D:1.2%] F:1.8% M:1.6% | C:86.2% [S:84.8%, D:1.4%] F:6.7% M:7.1% | C:94.7% [S:93.1%, D:1.6%] F:2.2% M:3.1% |
| Proteome BUSCO metazoa *odb10* (n = 954) | C:86.6% [S:85.3%, D:1.3%] F:6.3% M:7.1% | C:95.4% [S:92.0%, D:3.4%] F:2.3% M:2.3% | C:97.0% [S:95.8%, D:1.2%] F:1.5% M:1.5% | C:87.8% [S:86.3%, D:1.5%] F:5.5% M:6.7% | C:95.3% [S:93.7%, D:1.6%] F:1.7% M:3.0% |

**Supplemental Table S2.** RNA-sequencing datasets used for gene prediction.

| Species | SRA accession |
|---|---|
| *Cristatella mucedo* | SRR10620224 |
| *Cristatella mucedo* | SRR10620225 |
| *Cristatella mucedo* | SRR10620226 |
| *Cristatella mucedo* | SRR10620227 |
| *Cristatella mucedo* | SRR10620228 |
| *Cristatella mucedo* | SRR10620229 |
| *Cristatella mucedo* | SRR10620230 |
| *Cristatella mucedo* | SRR10620231 |
| *Watersipora subatra* | ERR11641139 |
| *Membranipora membranacea* | ERR6464929 |
| *Membranipora membranacea* | SRR2131259 |
| *Cryptosula pallasiana* | ERR9866430 |
| *Bugulina stolonifera* | ERR10123683 |
| *Bugulina stolonifera* | SRR11096622 |
| *Bugulina stolonifera* | SRR11096623 |
| *Bugulina stolonifera* | SRR11096624 |
| *Bugulina stolonifera* | SRR11096625 |
| *Bugulina stolonifera* | SRR11096626 |
| *Bugulina stolonifera* | SRR11096627 |
| *Bugulina stolonifera* | SRR11096628 |
| *Bugulina stolonifera* | SRR11096629 |
| *Bugulina stolonifera* | SRR11096631 |
| *Bugulina stolonifera* | SRR11096632 |
| *Bugulina stolonifera* | SRR11096632 |
| *Bugulina stolonifera* | SRR11096633 |
| *Bugulina stolonifera* | SRR11096634 |
| *Bugulina stolonifera* | SRR11096635 |
| *Bugulina stolonifera* | SRR11096636 |
| *Bugulina stolonifera* | SRR11096637 |
| *Bugulina stolonifera* | SRR11096638 |
| *Bugulina stolonifera* | SRR11096639 |
| *Bugulina stolonifera* | SRR11096640 |
| *Bugulina stolonifera* | SRR9667735 |

**Supplemental Table S3.** Repeat content of bryozoan genomes. Numbers correspond to the percentage of the genome composed by each type of element.

| | ***Bugulina stolonifera*** **(% of genome)** | ***Cristatella mucedo*** **(% of genome)** | ***Cryptosula pallasiana*** **(% of genome)** | ***Membranipora membranacea*** **(% of genome)** | ***Watersipora subatra*** **(% of genome)** |
|---|---|---|---|---|---|
| **Bases masked** | **32.15** | **47.52** | **57.01** | **35.5** | **53.21** |
| **Retroelements** | **9.31** | **25.19** | **20.91** | **8.99** | **9.43** |
| SINEs: | 0.28 | 0.02 | 0.08 | 0.36 | 0.14 |
| Penelope | 0.69 | 0.01 | 0 | 0 | 0.01 |
| LINEs | 4.01 | 23.96 | 3.64 | 5.01 | 4.18 |
| CRE/SLACS | 0 | 0 | 0 | 0.02 | 0 |
| L2/CR1/Rex | 2.99 | 23.22 | 2.51 | 2.91 | 1.63 |
| R1/LOA/Jockey | 0 | 0.07 | 0.02 | 0.13 | 0.02 |
| R2/R4/NeSL | 0.05 | 0.25 | 0.23 | 0.89 | 1.1 |
| RTE/Bov-B | 0.25 | 0.01 | 0.46 | 0.42 | 0.36 |
| L1/CIN4 | 0 | 0.1 | 0.15 | 0.26 | 0.52 |
| LTR | 5.01 | 1.2 | 17.19 | 3.62 | 5.1 |
| BEL/Pao | 0.45 | 0 | 0.25 | 0.39 | 0.44 |
| Ty1/Copia | 0.24 | 0.11 | 0.22 | 0.34 | 0.37 |
| Gypsy/DIRS1 | 2.9 | 1.01 | 12.77 | 1.76 | 4.03 |
| Retroviral | 0.15 | 0.08 | 0.13 | 0.22 | 0.23 |
| DNA transposons | 4.3 | 7.03 | 9.03 | 3.04 | 11.37 |
| hobo-Activator | 1.09 | 1.25 | 1.87 | 0.36 | 1.53 |
| Tc1-IS630-Pogo | 0.2 | 1.64 | 0.13 | 0.29 | 5.21 |
| En-Spm | 0 | 0 | 0 | 0 | 0 |
| MULE-MuDR | 0.09 | 0.26 | 0.21 | 0.28 | 0.19 |
| PiggyBac | 0.02 | 0 | 0.03 | 0.07 | 0.32 |
| Tourist/Harbinger | 0.11 | 0.02 | 0.17 | 0.21 | 0.07 |
| Other (Mirage, P-element, Transib) | 0.06 | 0.02 | 0.04 | 0.26 | 0.3 |
| **Rolling Circles** | **0.03** | **0.1** | **0.24** | **0.1** | **1.24** |
| **Unclassified** | **15.9** | **12.09** | **25.33** | **22.5** | **29.8** |
| **Satellites** | **0.17** | **0.09** | **0.03** | **0.03** | **0.17** |
| **Simple Repeats** | **1.68** | **2.68** | **1.4** | **0.76** | **1.09** |
| **Low complexity repeats** | **0.07** | **0.33** | **0.06** | **0.05** | **0.09** |

**Supplemental Table S4.** Genomes used for principal component analysis (PCA). Bryozoan species are in bold.

| Taxon | Species | Species ID | Assembly source | Assembly accession | Annotation source |
|---|---|---|---|---|---|
| Porifera | *Amphimedon queenslandica* | Aqu | NCBI | GCF_000090795.2 | NCBI RefSeq |
| Echinodermata | *Asterias rubens* | Aru | NCBI | GCF_902459465.1 | NCBI RefSeq |
| Rotifera | *Adineta vaga* | Ava | NCBI | GCA_021613535.1 | NCBI GenBank |
| Chordata | *Branchiostoma lanceolatum* | Bla | NCBI | GCA_927797965.1 | NCBI GenBank |
| Ctenophora | *Bolinopsis microptera* | Bmi | NCBI | GCA_026151205.1 | This work |
| **Bryozoa** | ***Bugulina stolonifera*** | **Bst** | **NCBI** | **GCA_935421135.1** | **This work** |
| Nematoda | *Caenorhabditis elegans* | Cel | NCBI | **GCA_000002985.3** | NCBI RefSeq |
| Chordata | *Ciona intestinalis* | Cin | NCBI | GCF_000224145.3 | NCBI RefSeq |
| **Bryozoa** | ***Cristatella mucedo*** | **Cmu** | **GEO** | **GSE169088** | **This work** |
| **Bryozoa** | ***Cryptosula pallasiana*** | **Cpa** | **NCBI** | **GCA_945261195.1** | **This work** |
| Annelida | *Capitella teleta* | Cte | NCBI | GCA_000328365.1 | NCBI GenBank |
| Arthropoda | *Drosophila melanogaster* | Dme | NCBI | GCF_000001215.4 | NCBI RefSeq |
| Arthropoda | *Daphnia pulex* | Dpu | NCBI | GCF_021134715.1 | NCBI RefSeq |
| Chordata | *Danio rerio* | Dre | NCBI | GCF_000002035.6 | NCBI RefSeq |
| Cnidaria | *Galaxea fascicularis* | Gfa | NCBI | GCA_948470475.1 | This work |
| Xenacoelomorpha | *Hofstenia miamia* | Hmi | Ensembl Metazoa | GCA_004352715.1 | Ensembl |
| Annelida | *Helobdella robusta* | Hro | NCBI | GCF_000326865.2 | NCBI RefSeq |
| Chordata | *Homo sapiens* | Hsa | NCBI | GCF_000001405.40 | NCBI RefSeq |
| Cnidaria | *Hydra vulgaris* | Hvu | NCBI | GCF_022113875.1 | NCBI RefSeq |
| Brachiopoda | *Lingula anatina* | Lan | Unpublish ed | Unpublished | This work |
| Nemertea | *Lineus longissimus* | Llo | NCBI | GCA_910592395.2 | This work |
| Echinodermata | *Lytechinus variegatus* | Lva | NCBI | GCF_018143015.1 | NCBI RefSeq |
| **Bryozoa** | ***Membranipora membranacea*** | **Mme** | **NCBI** | **GCA_914767715.1** | **This work** |
| Nemertea | *Notospermus geniculatus* | Nge | OIST | nge_genome_v2.0.fa.gz | OIST |
| Cnidaria | *Nematostella vectensis* | Nve | NCBI | GCF_932526225.1 | NCBI RefSeq |
| Mollusca | *Octopus bimaculoides* | Obi | NCBI | GCF_001194135.2 | NCBI RefSeq |
| Annelida | *Owenia fusiformis* | Ofu | NCBI | GCA_903813345.2 | NCBI RefSeq |
| Phoronida | *Phoronis australis* | Pau | OIST | pau_genome_v2.0.fa.gz | OIST |

| | | | | | |
|---|---|---|---|---|---|
| Mollusca | *Pecten maximus* | Pma | NCBI | GCF_902652985.1 | NCBI RefSeq |
| Mollusca | *Patella vulgata* | Pvu | NCBI | GCF_932274485.2 | NCBI RefSeq |
| Hemichordata | *Saccoglossus kowalevskii* | Sko | NCBI | GCF_000003605.2 | NCBI RefSeq |
| Platyhelminthes | *Schistosoma mansoni* | Sma | NCBI | GCF_000237925.1 | NCBI RefSeq |
| Platyhelminthes | *Schmidtea mediterranea* | Sme | SmedGD | SmedSxl_genome_v4.0 | SmedGD |
| Xenacoelomorpha | *Symsagittifera roscoffensis* | Sro | http://gb.macgenome.org | N/A | http://gb.macgenome.org |
| Placozoa | *Trichoplax adhaerens* | Tad | NCBI | GCF_000150275.1 | NCBI RefSeq |
| Arthropoda | *Tribolium castaneum* | Tca | NCBI | GCF_000002335.3 | NCBI RefSeq |
| **Bryozoa** | ***Watersipora subatra*** | **Wsu** | **NCBI** | **GCA_963576615.1** | **This work** |
| Chordata | *Xenopus tropicalis* | Xtr | NCBI | GCF_000004195.4 | NCBI RefSeq |

**Supplemental Table S5.** OrthoFinder output for bryozoan species.

| | |
|---|---|
| Number of species | 5 |
| Number of genes | 127,707 |
| Number of genes in orthogroups | 97,475 |
| Number of unassigned genes | 30,232 |
| Percentage of genes in orthogroups | 76.3 |
| Percentage of unassigned genes | 23.7 |
| Number of orthogroups | 16,984 |
| Number of species-specific orthogroups | 4621 |
| Number of genes in species-specific orthogroups | 22,441 |
| Percentage of genes in species-specific orthogroups | 17.6 |
| Mean orthogroup size | 5.7 |
| Median orthogroup size | 5 |
| G50 (assigned genes) | 6 |
| G50 (all genes) | 5 |
| O50 (assigned genes) | 4,544 |
| O50 (all genes) | 7,483 |
| Number of orthogroups with all species present | 6,765 |
| Number of single-copy orthogroups | 4,242 |

**Supplemental Table S6.** Pairwise microsynteny mixing scores for entire bryozoan genomes.

| Species 1 | Species 2 | Mixing score |
|---|---|---|
| CMU | MME | 0.551 |
| CMU | BST | 0.553 |
| CMU | WSU | 0.552 |
| CMU | CPA | 0.554 |
| MME | CPA | 0.046 |
| MME | WSU | 0.046 |
| MME | BST | 0.043 |
| BST | CPA | 0.019 |
| BST | WSU | 0.024 |
| WSU | CPA | 0.013 |

**Supplemental Table S7.** Pairwise microsynteny mixing scores for individual gymnolaemate bryozoan chromosomes.

| Species 1 | Species 2 | Species 1 chromosome | Species 2 chromosome | Order on dot plots | Mixing Rate | Estimated divergence time (MYA) |
|---|---|---|---|---|---|---|
| CPA | BST | 10 | 1 | 1 | 0.281 | 128 |
| CPA | BST | 12 | 1 | 2 | 0.8 | 128 |
| CPA | BST | 4 | 2 | 3 | 0.111 | 128 |
| CPA | BST | 6 | 3 | 4 | 0.634 | 128 |
| CPA | BST | 8 | 4 | 5 | 0.363 | 128 |
| CPA | BST | 5 | 11 | 6 | 0.407 | 128 |
| CPA | BST | 2 | 8 | 7 | 0.349 | 128 |
| CPA | BST | 1 | 5 | 8 | 0.343 | 128 |
| CPA | BST | 11 | 7 | 9 | 0.514 | 128 |
| CPA | BST | 7 | 9 | 10 | 0.732 | 128 |
| CPA | BST | 3 | 10 | 11 | 0.612 | 128 |
| CPA | BST | 9 | 6 | 12 | 0.638 | 128 |
| CPA | MME | 10 | 1 | 1 | 0.902 | 256 |
| CPA | MME | 12 | 1 | 2 | 0.667 | 256 |
| CPA | MME | 4 | 4 | 3 | 0.952 | 256 |
| CPA | MME | 6 | 2 | 4 | 0.914 | 256 |
| CPA | MME | 8 | 3 | 5 | 0.906 | 256 |
| CPA | MME | 5 | 7 | 6 | 0.715 | 256 |
| CPA | MME | 2 | 8 | 7 | 0.817 | 256 |
| CPA | MME | 1 | 6 | 8 | 0.968 | 256 |
| CPA | MME | 11 | 9 | 9 | 0.968 | 256 |
| CPA | MME | 7 | 10 | 10 | 0.856 | 256 |
| CPA | MME | 3 | 5 | 11 | 0.604 | 256 |
| CPA | MME | 9 | 11 | 12 | 0.926 | 256 |
| CPA | WSU | 10 | 1 | 1 | 0.134 | 64 |
| CPA | WSU | 12 | 1 | 2 | 0.689 | 64 |
| CPA | WSU | 4 | 3 | 3 | 0.099 | 64 |
| CPA | WSU | 6 | 4 | 4 | 0.528 | 64 |
| CPA | WSU | 8 | 2 | 5 | 0.081 | 64 |
| CPA | WSU | 5 | 6 | 6 | 0.062 | 64 |
| CPA | WSU | 2 | 5 | 7 | 0.02 | 64 |
| CPA | WSU | 1 | 7 | 8 | 0.458 | 64 |
| CPA | WSU | 11 | 9 | 9 | 0.499 | 64 |
| CPA | WSU | 7 | 11 | 10 | 0.42 | 64 |
| CPA | WSU | 3 | 8 | 11 | 0.279 | 64 |

| | | | | | | |
|---|---|---|---|---|---|---|
| CPA | WSU | 9 | 10 | 12 | 0.252 | 64 |
| MME | BST | 1 | 1 | 1 | 0.9515372 | 256 |
| MME | BST | 4 | 2 | 2 | 0.943304013 | 256 |
| MME | BST | 2 | 3 | 3 | 0.853222734 | 256 |
| MME | BST | 3 | 4 | 4 | 0.96521043 | 256 |
| MME | BST | 7 | 11 | 5 | 0.700711183 | 256 |
| MME | BST | 8 | 8 | 6 | 0.946323022 | 256 |
| MME | BST | 6 | 5 | 7 | 0.794550167 | 256 |
| MME | BST | 9 | 7 | 8 | 0.82755028 | 256 |
| MME | BST | 10 | 9 | 9 | 0.934399245 | 256 |
| MME | BST | 5 | 10 | 10 | 0.964606918 | 256 |
| MME | BST | 11 | 6 | 11 | 0.999654756 | 256 |
| MME | WSU | 1 | 1 | 1 | 0.9515372 | 256 |
| MME | WSU | 4 | 3 | 2 | 0.943304013 | 256 |
| MME | WSU | 2 | 4 | 3 | 0.853222734 | 256 |
| MME | WSU | 3 | 2 | 4 | 0.96521043 | 256 |
| MME | WSU | 7 | 6 | 5 | 0.700711183 | 256 |
| MME | WSU | 8 | 5 | 6 | 0.946323022 | 256 |
| MME | WSU | 6 | 7 | 7 | 0.794550167 | 256 |
| MME | WSU | 9 | 9 | 8 | 0.82755028 | 256 |
| MME | WSU | 10 | 11 | 9 | 0.934399245 | 256 |
| MME | WSU | 5 | 8 | 10 | 0.964606918 | 256 |
| MME | WSU | 11 | 10 | 11 | 0.999654756 | 256 |
| BST | WSU | 1 | 1 | 1 | 0.161199763 | 128 |
| BST | WSU | 2 | 3 | 2 | 0.211119054 | 128 |
| BST | WSU | 3 | 4 | 3 | 0.930367893 | 128 |
| BST | WSU | 4 | 2 | 4 | 0.472108731 | 128 |
| BST | WSU | 11 | 6 | 5 | 0.621155978 | 128 |
| BST | WSU | 8 | 5 | 6 | 0.522689494 | 128 |
| BST | WSU | 5 | 7 | 7 | 0.961633944 | 128 |
| BST | WSU | 7 | 9 | 8 | 0.384844331 | 128 |
| BST | WSU | 9 | 11 | 9 | 0.933948334 | 128 |
| BST | WSU | 10 | 8 | 10 | 0.680422252 | 128 |
| BST | WSU | 6 | 10 | 11 | 0.409792086 | 128 |

**Supplemental Table S8.** ALG composition of bilaterian genomes.

| **Phylum** | **Species** | **Chr.** | **Bilaterian ALGs** |
|---|---|---|---|
| Chordata | *B. floridae* | 1 | A1, A2 |
| | | 2 | C1, J2 |
| | | 3 | C2, Q |
| | | 4 | I, O1 |
| | | 5 | E |
| | | 6 | D |
| | | 7 | F |
| | | 8 | M |
| | | 9 | K |
| | | 10 | B1 |
| | | 11 | G |
| | | 12 | N |
| | | 13 | H |
| | | 14 | P |
| | | 15 | L |
| | | 16 | B2 |
| | | 17 | J1 |
| | | 18 | B3 |
| | | 19 | O2 |
| Mollusca | *P. maximus* | 1 | H, Q |
| | | 2 | B2, M |
| | | 3 | A1 |
| | | 4 | J2, L |
| | | 5 | K, O2 |
| | | 6 | F |
| | | 7 | E |
| | | 8 | G |
| | | 9 | C1 |
| | | 10 | O1, R |
| | | 11 | D |
| | | 12 | I |
| | | 13 | A2 |
| | | 14 | N |
| | | 15 | B1 |
| | | 16 | P |
| | | 17 | C2 |

| | | | |
|---|---|---|---|
| | | 18 | J1 |
| | | 19 | B3 |
| Nemertea | *L. longissimus* | 1 | H, Q |
| | | 2 | C1, G |
| | | 3 | A1 |
| | | 4 | J2, L |
| | | 5 | K, O2 |
| | | 6 | I |
| | | 7 | E |
| | | 8 | O1, R |
| | | 9 | F |
| | | 10 | N |
| | | 11 | D |
| | | 12 | P |
| | | 13 | B2 |
| | | 14 | C2 |
| | | 15 | B3 |
| | | 16 | B1 |
| | | 17 | M |
| | | 18 | A2 |
| | | 19 | J1 |
| Brachiopoda | *L. anatina* | 1 | C1, G, I, J2, K, L, O1, O2, R |
| | | 2 | A1, B3, C2, H, Q |
| | | 3 | F, M |
| | | 4 | D |
| | | 5 | B1 |
| | | 6 | E |
| | | 7 | A2, B2 |
| | | 8 | J1 |
| | | 9 | P |
| | | 10 | N |

**Supplemental Table S9.** ALG composition of bryozoan genomes.

| Species | Chr. | Bilaterian ALGs | Bryozoan ALGs | Gymnolaemata ALGs |
|---|---|---|---|---|
| CMU | 1 | A1, A2, B1, B2, B3, C1, D, E, G, H, I, J1, J2, L, O1, Q, R, M | α, β, γ | *a, b, c, d, e, f, g, h, i, j, k* |
| | 2 | A1, A2, B1, B2, B3, C1, D, E, F, G, H, I, J1, J2, L, N, O1, Q, R, (M) | α, β, ζ | *a, b, c, d, g, h, j, k* |
| | 3 | A1, B1, B2, B3, C1, D, E, G, K, L, O2, P | β, ε | *a, b, c, d, g, h, i, j, k* |
| | 4 | A1, B1, B2, B3, C1, C2, D, E, F, G, K, L, N, O2, P | β, δ, ε, ζ | *a, b, c, d, f, g, h, i, j, k* |
| | 5 | A2, H, I , J1, J2, L, M O1, Q, R | α, γ | *a, b, c, d, e, f, g, h, i, k* |
| | 6 | A1, F, N | δ, ζ | *a, b, c, f, g, j, k* |
| | 7 | A1, B1, B2, B3, C1, D, E, G, K, L, O2, P | β, ε | *a, b, c, d, h, i* |
| | 8 | A1, C2 | δ | *f, g* |
| MME | 1 | A1, A2, B1, B2, B3, C1, D, E, G, H, I, J1, J2, L, O1, Q, R, (M) | α, β, (γ) | *a* |
| | 2 | A1, A2, B1, B2, B3, C1, D, E, G, H, I, J1, J2, L, O1, Q, R, (M) | α, β, (γ) | *c* |
| | 3 | A1, A2, B1, B2, B3, C1, D, E, G, H, I, J1, J2, L, O1, Q, R, (M) | α, β, (γ) | *d* |
| | 4 | A1, A2, B1, B2, B3, C1, D, E, G, H, I, J1, J2, L, O1, Q, R, (M) | α, β, (γ) | *b* |
| | 5 | F, N, (M) | ζ, (γ) | *j* |
| | 6 | A1, C2, (M) | δ, (γ) | *g* |
| | 7 | M | γ | *e* |
| | 8 | A1, C2, (M) | δ, (γ) | *f* |
| | 9 | K, L, O2, P, (M) | ε, (γ) | *h* |
| | 10 | K, L, O2, P, (M) | ε, (γ) | *i* |
| | 11 | F, N, (M) | ζ, (γ) | *k* |
| BST | 1 | A1, A2, B1, B2, B3, C1, D, E, G, H, I, J1, J2, L, O1, Q, R, (M) | α, β, (γ) | *a* |
| | 2 | A1, A2, B1, B2, B3, C1, D, E, G, H, I, J1, J2, L, O1, Q, R, (M) | α, β, (γ) | *b* |
| | 3 | A1, A2, B1, B2, B3, C1, D, E, G, H, I, J1, J2, L, O1, Q, R, (M) | α, β, (γ) | *c* |
| | 4 | A1, A2, B1, B2, B3, C1, D, E, G, H, I, J1, J2, L, O1, Q, R, (M) | α, β, (γ) | *d* |
| | 5 | A1, C2, (M) | δ, (γ) | *g* |
| | 6 | F, N, (M) | ζ, (γ) | *k* |
| | 7 | K, L, O2, P, (M) | ε, (γ) | *h* |
| | 8 | A1, C2, (M) | δ, (γ) | *f* |
| | 9 | K, L, O2, P, (M) | ε, (γ) | *i* |
| | 10 | F, N, (M) | ζ, (γ) | *j* |
| | 11 | M | γ | *e* |
| WSU | 1 | A1, A2, B1, B2, B3, C1, D, E, G, H, I, J1, J2, L, O1, Q, R, (M) | α, β, (γ) | *a* |
| | 2 | A1, A2, B1, B2, B3, C1, D, E, G, H, I, J1, J2, L, O1, Q, R, (M) | α, β, (γ) | *d* |

| | | | | |
|---|---|---|---|---|
| | 3 | A1, A2, B1, B2, B3, C1, D, E, G, H, I, J1, J2, L, O1, Q, R, (M) | α, β, (γ) | *b* |
| | 4 | A1, A2, B1, B2, B3, C1, D, E, G, H, I, J1, J2, L, O1, Q, R, (M) | α, β, (γ) | *c* |
| | 5 | A1, C2, (M) | δ, (γ) | *f* |
| | 6 | M | γ | *e* |
| | 7 | A1, C2, (M) | δ, (γ) | *g* |
| | 8 | F, N, (M) | ζ, (γ) | *j* |
| | 9 | K, L, O2, P, (M) | ε, (γ) | *h* |
| | 10 | K, L, O2, P, (M) | ε, (γ) | *k* |
| | 11 | F, N, (M) | ζ, (γ) | *i* |
| CPA | 1 | A1, C2, (M) | δ, (γ) | *g* |
| | 2 | A1, C2, (M) | δ, (γ) | *f* |
| | 3 | F, N, (M) | ζ, (γ) | *j* |
| | 4 | A1, A2, B1, B2, B3, C1, D, E, G, H, I, J1, J2, L, O1, Q, R, (M) | α, β, (γ) | *b* |
| | 5 | M | γ | *e* |
| | 6 | A1, A2, B1, B2, B3, C1, D, E, G, H, I, J1, J2, L, O1, Q, R, (M) | α, β, (γ) | *c* |
| | 7 | K, L, O2, P, (M) | ε, (γ) | *i* |
| | 8 | A1, A2, B1, B2, B3, C1, D, E, G, H, I, J1, J2, L, O1, Q, R, (M) | α, β, (γ) | *d* |
| | 9 | F, N, (M) | ζ, (γ) | *k* |
| | 10 | A1, A2, B1, B2, B3, C1, D, E, G, H, I, J1, J2, L, O1, Q, R, (M) | α, β, (γ) | *a* |
| | 11 | K, L, O2, P, (M) | ε, (γ) | *h* |
| | 12 | A1, A2, B1, B2, B3, C1, D, E, G, H, I, J1, J2, L, O1, Q, R, (M) | α, β, (γ) | *a* |

**Supplemental Table S10.** Origins of genes in bryozoan genomes with the annelid *Metaphire vulgaris* for comparison.

| | ***Metaphire vulgaris* (known WGD)** | ***Membranipora membranacea*** | ***Cryptosula pallasiana*** | ***Bugulina stolonifera*** | ***Watersipora subatra*** | ***Cristatella mucedo*** |
|---|---|---|---|---|---|---|
| Singleton | 43.0 | 39.8 | 36.1 | 44.7 | 40.1 | 68.6 |
| Dispersed duplicate | 25.0 | 33.4 | 34.0 | 32.8 | 34.9 | 17.7 |
| Proximal duplicate | 3.9 | 12.2 | 13.6 | 7.8 | 11.0 | 6.6 |
| Tandem duplicate | 4.6 | 13.1 | 14.6 | 14.3 | 10.4 | 5.0 |
| WGD/segmental duplicate | 23.5 | 1.5 | 1.7 | 0.4 | 3.6 | 2.1 |

**Supplemental Table S11.** Composition of bryozoan ALGs.

| Bryozoan ALG | Bilaterian ALGs |
|---|---|
| α | A2, H, I, J1, J2, L, O1, Q, R, M |
| β | B1, B2, B3, C1, D, E, G, A1 |
| γ | M |
| δ | A1, C2 |
| ε | K, O2, P, L |
| ζ | F, N |

**Supplemental Table S12.** Summary statistics for the *Lingula anatina* genome assembly.

| **Description** | ***Lingula anatina* statistic** |
|---|---|
| Genome size (Mb) | 329 |
| Genome size (chromosome-level scaffolds only, Mb) | 322 |
| Sequencing coverage | 405-fold |
| Number of scaffolds | 16 |
| Number of scaffolds > 10 Mb | 10 |
| Scaffold N50 (Mb) | 30.1 |
| Contig N50 (kb) | 22,282 |
| GC content (%) | 36.5 |
| Repeats (%) | 31.7 |
| Genome BUSCO metazoa *odb10* (n = 954) | C:97.5%[S:96.6%,D:0.9%],F:1.8%,M:0.7% |

**Supplemental Table S13.** Hi-C scaffolding statistics for the *Lingula anatina* genome assembly.

| Superscaffold | Number of contings | Length of contigs | Length of superscaffold |
|---|---|---|---|
| chr1 | 6 | 71,918,473 | 71,920,973 |
| chr2 | 2 | 49,366,815 | 49,367,315 |
| chr3 | 1 | 34,252,355 | 34,252,355 |
| chr4 | 4 | 30,084,556 | 30,086,056 |
| chr5 | 6 | 25,661,712 | 25,664,212 |
| chr6 | 2 | 24,769,930 | 24,770,430 |
| chr7 | 3 | 24,364,059 | 24,365,059 |
| chr8 | 3 | 20,822,434 | 20,823,434 |
| chr9 | 4 | 20,476,484 | 20,477,984 |
| chr10 | 3 | 20,310,182 | 20,311,182 |
| TOTAL | 34 | 322,027,000 | 322,039,000 |

**Supplemental Table S14.** Expression levels of bryozoan Hox genes in RNA-seq datasets. TPM = transcripts per [kilobase] million.

| Species | Tissue | SRA accession | Transcript | Gene | TPM |
|---|---|---|---|---|---|
| CMU | Adult | SRR10620224 | g8621.t1 | *Hox4a* | 0.692 |
| CMU | Adult | SRR10620224 | g5048.t1 | *Hox4b* | 0.966 |
| CMU | Adult | SRR10620224 | g8615.t1 | *Lox5a* | 0.611 |
| CMU | Adult | SRR10620224 | g5021.t1 | *Lox5b* | 5.712 |
| CMU | Adult | SRR10620225 | g8621.t1 | *Hox4a* | 0.000 |
| CMU | Adult | SRR10620225 | g5048.t1 | *Hox4b* | 0.964 |
| CMU | Adult | SRR10620225 | g8615.t1 | *Lox5a* | 0.445 |
| CMU | Adult | SRR10620225 | g5021.t1 | *Lox5b* | 3.563 |
| CMU | Adult | SRR10620226 | g8621.t1 | *Hox4a* | 0.487 |
| CMU | Adult | SRR10620226 | g5048.t1 | *Hox4b* | 1.606 |
| CMU | Adult | SRR10620226 | g8615.t1 | *Lox5a* | 0.233 |
| CMU | Adult | SRR10620226 | g5021.t1 | *Lox5b* | 5.139 |
| CMU | Adult | SRR10620227 | g8621.t1 | *Hox4a* | 0.374 |
| CMU | Adult | SRR10620227 | g5048.t1 | *Hox4b* | 1.890 |
| CMU | Adult | SRR10620227 | g8615.t1 | *Lox5a* | 0.630 |
| CMU | Adult | SRR10620227 | g5021.t1 | *Lox5b* | 5.518 |
| CMU | Adult | SRR10620228 | g8621.t1 | *Hox4a* | 0.146 |
| CMU | Adult | SRR10620228 | g5048.t1 | *Hox4b* | 1.194 |
| CMU | Adult | SRR10620228 | g8615.t1 | *Lox5a* | 0.907 |
| CMU | Adult | SRR10620228 | g5021.t1 | *Lox5b* | 5.874 |
| CMU | Adult | SRR10620229 | g8621.t1 | *Hox4a* | 0.478 |
| CMU | Adult | SRR10620229 | g5048.t1 | *Hox4b* | 1.076 |
| CMU | Adult | SRR10620229 | g8615.t1 | *Lox5a* | 0.336 |
| CMU | Adult | SRR10620229 | g5021.t1 | *Lox5b* | 4.276 |
| CMU | Adult | SRR10620230 | g8621.t1 | *Hox4a* | 0.602 |
| CMU | Adult | SRR10620230 | g5048.t1 | *Hox4b* | 0.978 |
| CMU | Adult | SRR10620230 | g8615.t1 | *Lox5a* | 0.116 |
| CMU | Adult | SRR10620230 | g5021.t1 | *Lox5b* | 3.529 |
| CMU | Adult | SRR10620231 | g8621.t1 | *Hox4a* | 0.498 |
| CMU | Adult | SRR10620231 | g5048.t1 | *Hox4b* | 1.106 |
| CMU | Adult | SRR10620231 | g8615.t1 | *Lox5a* | 0.236 |
| CMU | Adult | SRR10620231 | g5021.t1 | *Lox5b* | 2.681 |

**Legends for Supplemental Datasets**

**Supplemental Data S1 (separate file).** Table of single-copy orthologs used in this study with their corresponding ALGs. Column explanations: (1) orthogroup ID number from OrthoFinder; (2) bilaterian ALG to which genes belong (4 – 9) Gene IDs for each species.

**Supplemental Data S2 (separate file).** Table of homeodomain sequences for Hox genes annotated in this work. Column explanations: (1) species name; (2) gene name; (3) alternate gene name; (4) chromosome location of gene – chromosome number; (5) chromosome location of gene – chromosome accession; (6) strand of gene; (7) start position of homeodomain exon 1; (8) end position of homeodomain exon 1; (9) start position of homeodomain exon 2; (10) end position of homeodomain exon 2; (11) amino acid sequence of homeodomain.
